# Rapid Kinetic Fingerprinting of Single Nucleic Acid Molecules by a FRET-based Dynamic Nanosensor

**DOI:** 10.1101/2021.04.06.438627

**Authors:** Kunal Khanna, Shankar Mandal, Aaron T. Blanchard, Muneesh Tewari, Alexander Johnson-Buck, Nils G. Walter

## Abstract

Biofluids contain cell-free nucleic acids such as microRNAs (miRNAs) and circulating tumor-derived DNAs (ctDNAs) that have emerged as promising disease biomarkers. Conventional detection of these biomarkers by digital PCR and next generation sequencing, although highly sensitive, requires time-consuming extraction and amplification steps that increase the risk of sample loss and cross-contamination, respectively. To achieve the direct, rapid detection of miRNAs and ctDNAs with near-perfect specificity and single-molecule level sensitivity, we herein describe an accelerated amplification-free single-molecule kinetic fingerprinting. This approach, termed intramolecular single-molecule recognition through equilibrium Poisson sampling (iSiMREPS), exploits a dynamic DNA nanosensor comprising a surface anchor and a pair of fluorescent detection probes: one probe captures individual target molecules onto the surface, while the other transiently interrogates them to generate kinetic fingerprints by intramolecular single-molecule Förster resonance energy transfer (smFRET). Formamide is used to further accelerate the kinetics of probe-target interactions and fingerprinting, while background signals are reduced by removing non-target-bound probes from the surface using toehold-mediated strand displacement. We show that iSiMREPS can detect in as little as 10 seconds two distinct, promising cancer biomarkers—miR-141 and a common *EGFR* exon 19 deletion—reaching a limit of detection (LOD) of ~3 fM and a mutant allele fraction among excess wild-type as low as 1 in 1 million, or 0.0001%. We anticipate that iSiMREPS will find utility in research and clinical diagnostics based on its features of rapid detection, high specificity, sensitivity, and generalizability.

## INTRODUCTION

Circulating cell-free nucleic acids (cfNAs) have emerged as promising diagnostic biomarkers for the detection of diseases such as cancer^1, 2^. Among various cfNAs, microRNAs (miRNAs) are short non-coding RNAs with gene regulatory function and great potential as biomarkers. Because miRNAs are generally associated with proteins, they are protected them from RNase degradation and, thus, occur at relatively high concentrations in the biofluids of cancer patients^2^. Another important class of cfNAs entails circulating, cell-free tumor-derived DNAs (ctDNAs), such as mutant copies of the epidermal growth factor receptor (*EGFR*) gene that are commonly found in the blood of some patients with non-small cell lung cancer (NSCLC)^3^. Detecting abnormal levels of these biomarkers in biofluids (e.g., blood, urine) via non-invasive liquid biopsies in place of traditional invasive tissue biopsies has been a major area of clinical interest^4, 5^. Monitoring of these cfNAs for diagnosis, prognosis and treatment evaluation has thus become increasingly important^2^ and has necessitated cfNA detection approaches that are rapid, highly specific, and highly sensitive.

cfNA detection from biofluids, however, can pose some technical challenges. ctDNA mutants, for example, often have very low allelic frequency (e.g., 0.1-1%)^6^ and must be measured against the background of the far more abundant wild-type DNA from normal cells. Technical challenges are also faced for miRNA detection, which typically requires complex, laborious procedures that include processing of sample matrix, purification, adapter ligation, and reverse transcription^7, 8^. Techniques such as next generation sequencing (NGS) for large-scale genome analysis^9^ and digital PCR^10^ have arisen as gold standards for nucleic acid detection. While they achieve high sensitivity for low abundance analytes and have sufficient specificity for allelic frequencies as low as 0.01% for ctDNA mutants, they require significant sample preparation, enzymatic reactions, and amplification steps that can introduce various errors and compromise assay performance when high specificity is necessary^11^. While several amplification-free methods^12–14^ for detecting cfNAs have been reported, the clinical utility of these techniques is constrained by upper limits on specificity imposed by the thermodynamics of nucleic acid binding^15^.

We recently reported an amplification-free single-molecule kinetic fingerprinting technique called Single Molecule Recognition through Equilibrium Poisson Sampling (SiMREPS)^16, 17^ that permits the ultraspecific and highly sensitive detection of miRNAs and ctDNAs from biofluids. SiMREPS uses single-molecule fluorescence microscopy to record the transient binding and dissociation of fluorescent probes to a surface-captured nucleic acid. Continuous imaging for ~10 minutes or longer reveals repeated binding to individual captured molecules, yielding a time-resolved “kinetic fingerprint” that can be used to distinguish a target molecule from non-target molecules and non-specific background^16, 17^. Imaging multiple fields-of-view (FOV_s_) can improve the sensitivity by increasing the odds of observing low-abundance target molecules, but adds further time needed for testing a given sample. As such, the time taken to collect data limits the full potential of SiMREPS as a rapid, single molecule nucleic acid assay technology, especially for high testing-volume clinical settings where hundreds of samples per day may need to be analysed.

Thus, shorter data acquisition times per FOV would improve the speed, sensitivity, and throughput of SiMREPS. Since the high specificity of SiMREPS requires the observation of multiple (>10) probe binding events to each molecule, an obvious strategy for accelerating data acquisition would be to increase the fluorescent probe concentration so that probe binding events are observed with higher frequency. However, the background fluorescence from unbound probes diffusing near the surface imposes an upper limit on the concentration of probe that can be used while maintaining sufficient signal-to-noise for single-molecule detection. Subsequently, this limits the maximum probe binding frequency that can be achieved in conventional SiMREPS.

To reduce signal from unbound fluorescent probe while increasing its local concentration, we developed a single-molecule Förster resonance energy transfer (smFRET)-based intramolecular kinetic fingerprinting approach termed intramolecular SiMREPS, or iSiMREPS (**Figure 1**). iSiMREPS introduces a dynamic DNA nanoscale sensor comprising an immobilized anchor stably hybridized to a pair of fluorescent capture and query probes. Transient intramolecular interactions within the sensor result in rapid transitions between high- and low-FRET states in the presence of the correct target molecule, while showing almost no high-FRET signal in the absence of the target or in the presence of related non-target sequences (**Figure 1**). These FRET transitions reveal a characteristic kinetic fingerprint of the analyte that reduces false positives dramatically compared to a static readout, since the signal must not only satisfy intensity thresholds but also exhibit a specific kinetic signature (**Figure 1**). The number of fluorescent probe binding and dissociation events (*N_b+d_*) to a single analyte molecule can be modeled as a Poisson process, with a coefficient of variation (C.V.) that decreases as *N_b+d_* increases. Consequently, the kinetic fingerprint from the analyte can be readily distinguished from non-specific binding to a near-perfect extent if the *N_b+d_* is large enough within the chosen observation time window, which can be extended as needed^16^. iSiMREPS allows for rapid stochastic transitions because the nanoscale intramolecular arrangement of the sensor creates a high local concentration of fluorescent probe (**Figure 1**), and thus significantly reduces the time between a dissociation event and the next binding event (*τ_off_*). The rate of the transitions between high-and low-FRET states can be further increased by reducing the thermodynamic stability of the probe-target complex (e.g. by minimizing complementary base pairs, modifying the lengths of the sensor strands to facilitate transitions between both states, increasing temperature, or adding denaturant). Here, we show that iSiMREPS can detect both a miRNA and a ctDNA with a limit of detection (LOD) of ~3 fM. The exon 19 deletion ctDNA is detected at mutant allele fractions as low as 1 in 1 million, corresponding to a specificity of >99.9999%. iSiMREPS acquisitions require approximately 10 s per FOV, an ~60-fold improvement compared to conventional (intermolecular) SiMREPS measurements that paves the way for accelerated molecular diagnostics.

**Figure 1.**
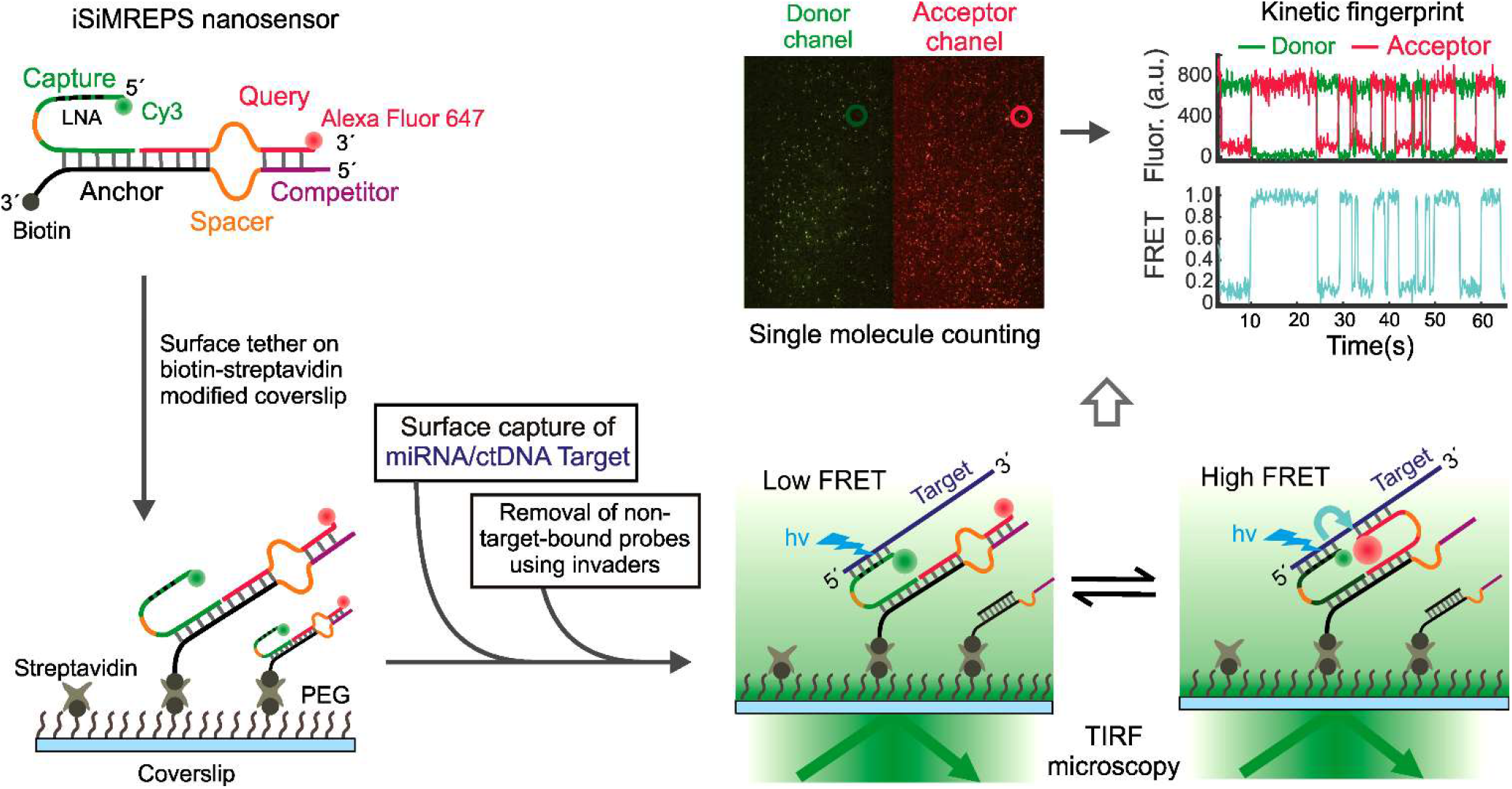
Schematic of iSiMREPS sensors for rapid kinetic fingerprinting of single nucleic acids. The iSiMREPS nanosensor consists of a biotinylated, surface-tethered anchor hybridized to a capture probe and a query probe. The capture probe is labeled with a donor fluorophore (Cy3) and partially modified with locked nucleic acid residues for high affinity capture of a miRNA/ctDNA target molecule. The query probe contains an acceptor fluorophore (Alexa Fluor 647) and transiently alternates between binding target and competitor sequences to generate high- and low-FRET signals, respectively. Transitions between FRET states are recorded by total internal reflection fluorescence (TIRF) microscopy, and FRET vs. time traces are analysed computationally to count single biomarker molecules with high specificity.

## RESULTS AND DISCUSSION

### The general architecture and working principles of smFRET-based iSiMREPS nanosensors for nucleic acids

smFRET-based iSiMREPS for counting single nucleic acid molecules (i.e., miRNA and ctDNA) utilizes a surface-immobilized nanoscale sensor composed of three DNA oligonucleotides: a biotinylated anchor (A), a donor (Cy3)-labeled capture probe, and an acceptor (Alexa Fluor 647, A647)-labeled query probe (**Figure 1**). The anchor stably binds the non-labeled ends of the capture and query probes via two 12 base pair (bp) duplexes of high GC content (~75% for capture sequence, ~83% for query sequence). The free end of the capture probe is partially modified with locked nucleic acid (LNA) residues, which enable high-affinity and kinetically stable capture of the miRNA/DNA target molecule. The free end of the query probe is designed to alternate between transient hybridization to the free end of the captured target nucleic acid molecule and a competitor (C) sequence that extends from the free end of the anchor. We introduced poly-deoxy-thymine (poly-dT) segments as spacers in the anchor, capture, and query strands to introduce the flexibility necessary for the sensor to properly assemble and transition between target-bound and competitor-bound states.

In the optimized assay, iSiMREPS sensors composed of pre-hybridized anchor, capture, and query strands are tethered first to a biotin-Bovine Serum Albumin (BSA)-passivated quartz slide (for prism-type total internal reflection fluorescence, or p-TIRF) or a PEG-passivated glass coverslip (for objective-type TIRF or o-TIRF) via streptavidin-biotin affinity linkages. The nucleic acid target molecules are then introduced into the solution above the surface. The target molecules bind strongly to the capture probes and thus become tethered to the surface. To minimize background signals before imaging with TIRF microscopy, non-target-bound fluorescent probes are removed from the surface by toehold mediated strand displacement (TMSD) using a pair of capture and query invader strands. As detailed below, this TMSD step proved important because cross-talk from the donor fluorophore into the acceptor detection channel, as well as a small amount of direct excitation of the acceptor fluorophore, result in high fluorescent background signal when sensors are present at the high surface densities required for efficient target capture.

In the presence of a target molecule, the query probe alternates between transiently binding to the target and the competitor, yielding distinct FRET signatures depending on which sequence is bound. When the query probe binds to the target, the donor and acceptor fluorophores are in close proximity, resulting in a high-FRET signal (**Figure 1**). In contrast, when the query probe dissociates from the target and/or binds to the competitor, the two fluorophores are far apart resulting in little to no FRET signal (**Figure 1**). The repeated transitions between high- and low-FRET signals generate a characteristic kinetic fingerprint, permitting the accurate identification of single target nucleic acid molecules. Because these transitions occur much more rapidly than the transitions in conventional (intermolecular) SiMREPS, we anticipated that smFRET-based iSiMREPS should allow for faster and higher-confidence detection of nucleic acids through rapid fingerprint generation. In the following sections, we optimize this general design to detect two distinct nucleic acid biomarkers of disease.

### Optimization of an iSiMREPS sensor design for detecting miRNA

An initial proof-of-concept iSiMREPS sensor was designed to detect the small non-coding RNA miR-141, a miRNA that has emerged as a biomarker for prostate cancer^18, 19^. The specific sensor design for detecting miR-141 is detailed in **Figure 2A**. To develop iSiMREPS into an accelerated single-molecule kinetic fingerprinting technique, we initially tested several sensor designs (**Figure S1A-C**) aiming to yield rapidly reversible smFRET transitions in the presence of target. To refer to different sensor designs, we use a Q_a_C_b_QS_c_CS_d_ naming convention, where Q is the query sequence complementary to target, QS is the query spacer, C is the competitor sequence complementary to the query sequence, CS is the competitor spacer, and the letters a, b, c, and d are integers reflecting the number of nucleotides in each domain (**Figure 2A**).

**Figure 2.**
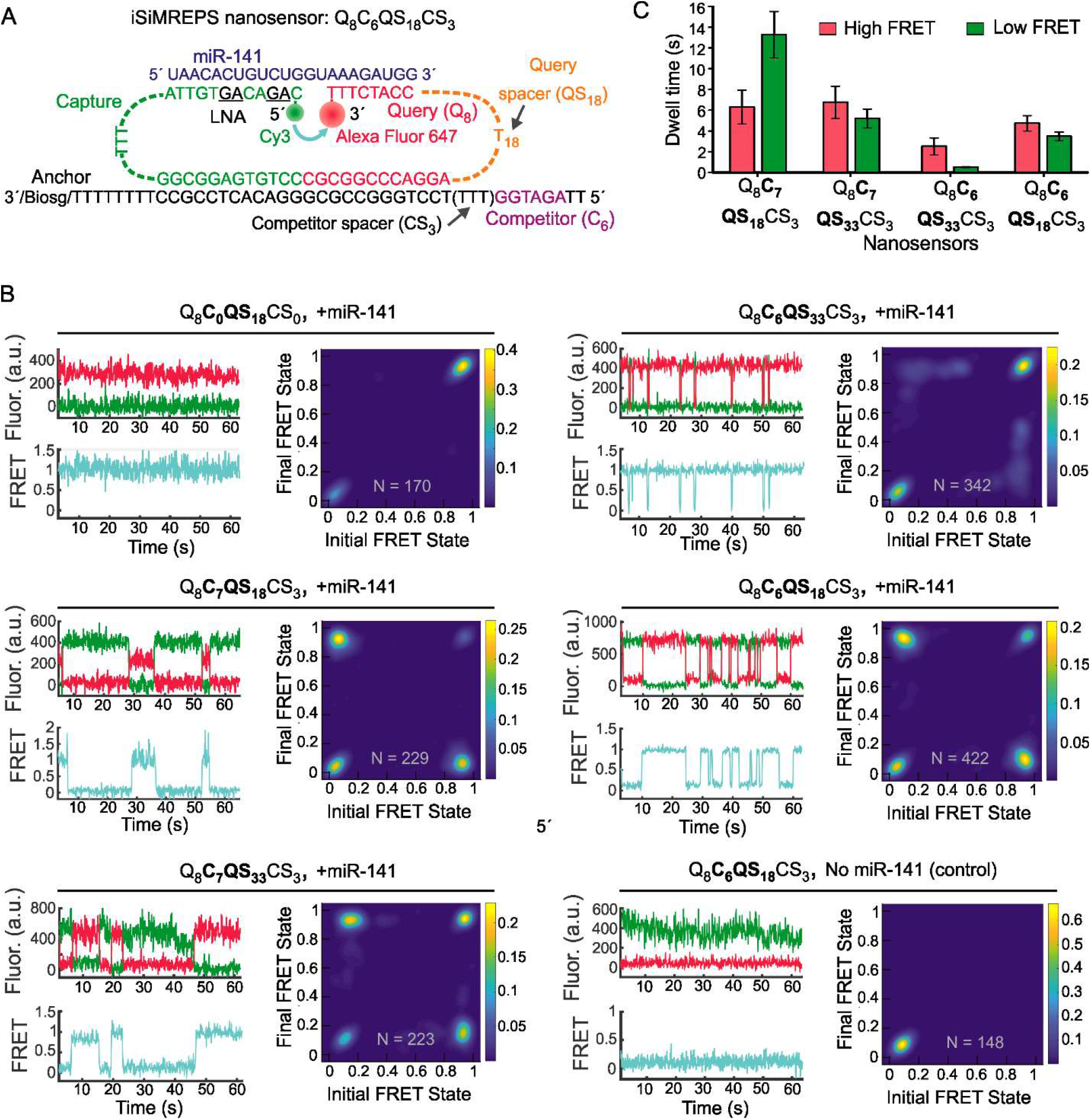
Design and optimization of iSiMREPS for detection of a miRNA. (A) Design of the optimized Q_8_C_6_QS_18_CS_3_ smFRET-based iSiMREPS sensor for detection of miR-141. The capture probe stably binds with the miRNA target with the assistance of locked nucleic acid residues (black and underlined) that increase the stability of the DNA-RNA duplex. The query probe (8 nt) switches between being bound to the 8 nt overhang of the target or to a 6 nt competitor sequence that extends from the anchor, resulting in dynamic kinetic smFRET fingerprints. (B) TODP plots and representative traces for different iSiMREPS sensor designs that have fixed query (8 nt), varying competitor (6 and 7 nt), fixed competitor spacer (3 nt), and varying query spacer (3, 18 and 33 nt) lengths in the presence of miR-141, as well as control without miR-141. The smFRET dynamics of each sensor is indicated. (C) The average lifetimes of the high-FRET (red) and low-FRET (green) interactions for each sensor design. Lifetime averages were calculated using an exponential decay fitting model, reporting averages calculated from either a single-exponential fit (if the sum squared error, sse, was < 0.05 and the R^2^ > 0.98) or a weighted double-exponential fit. All data are presented as mean ± s.d., where n = 3 populations of a split data set for each condition.

The stable and high affinity capture of miR-141 target molecule on slide surfaces was achieved by a 12 nucleotide (nt) segment of the capture probe with 4 locked nucleic acid (LNA)-modified residues that raised the *T_m_* of the capture probe-target duplex to 73°C. An 8 nt segment of the A647-tagged query probe (Q_8_) (*T_m_* = 30.2°C, see **Methods, and Table S1**) was designed to alternate between transient interactions with the target and competitor sequence to generate high-and low-FRET signals, respectively (**Figure 2A**). We initially omitted a competitor sequence (CS_0_) and instead used an 18-nt poly-dT (dT_18_) query spacer (QS_18_) to introduce conformational flexibility^20^ in the query strand and generate high-and low-FRET signals (**Figure S1A**). Prism-type TIRF characterization of the Q_8_C_0_QS_18_CS_0_ sensor in the presence of miR-141 (**see Methods, Figure S1A**) showed clear smFRET signals, suggesting that the sensor hybridized successfully with miR-141 to induce a high-FRET state (**Figure 2B, top left panel**). However, the equilibrium FRET distribution overwhelmingly favored the high-FRET state (**Figure 2B, top left panel**), preventing the characterization of a kinetic fingerprint. This heavy bias likely occurred due to the desired high local effective concentrations^21^ of the probes and target within the volume occupied by the assembled nanosensor, resulting in a high rate of query and target association. Consequently, while query-target dissociation events are expected, they appear too short-lived to be detected at the 60-100 ms time resolution achievable in smFRET. To disfavor the query-target interaction by increasing the entropic cost of hybridization, we increased the length of the query spacer from dT18 to dT33 (**Figure S1A**). However, this Q_8_C_0_QS_33_CS_0_ sensor still heavily favored the high-FRET state **(Figure S1D).**

We hypothesized that the addition of a competitor sequence could decrease the observed lifetime of the target bound (high-FRET) state by competing with the target to transiently bind to the query probe instead, thus stabilizing the non-target-bound (low-FRET) state and enabling the generation of dynamic smFRET signals (**Figure 1**). Therefore, in our next design we introduced a 7-nt competitor sequence (C_7_) (*T_m_*=18.1°C, **see Methods, and Table S1**) to the anchor and used a 3-nt polythymine (dT_3_) as a competitor spacer (CS_3_**) (Figure S1B).** The Q_8_C_7_QS_18_CS_3_ sensor showed repeated transitions between high-and low-FRET states that constituted a distinctive kinetic fingerprint in the presence of miR-141 (**Figure 2B, middle left panel).** In fact, this design was somewhat biased towards the low-FRET state. This experiment demonstrated that the competitor sequence is required for iSiMREPS designs to exhibit measurable transitions between smFRET states.

To estimate the lifetimes of smFRET states, each intensity-time trace was fit with a two-state Hidden Markov Model (HMM)^22^, and the dwell times of individual events were extracted (**Figure S2A**). For each of the two states, the average dwell time was then calculated by fitting an exponential decay function to the cumulative frequency of the dwell time population (**Figure S2B-S2E, and SI for detail**). These dwell times encompass the total lifetime spent in a particular FRET state before transitioning, and thus include events where the query probe may have instantly rebound to the same strand. Accordingly, the average dwell times reported here are best described as the mean first passage times between bound states (with the unbound state serving as a short-lived intermediate)^23^. The average dwell times for high- and low-FRET states for the Q_8_C_7_QS_18_CS_3_ sensor were 6.3 ± 1.6 s and 13.3 ± 2.2 s, respectively (**Figure 2C**). Interestingly, the average dwell time of the low-FRET state was approximately 2-fold higher than that of the high-FRET state (**Figure 2C**) even though the query-competitor duplex was less thermodynamically stable (*T_m_* = 18.1°C) than the query-target duplex (*T_m_* = 30.2°C, see **Table S1**), suggesting that the geometry of the sensor causes the query probe to preferentially bind the competitor rather than the target. Together, these results suggested that single-molecule kinetic fingerprinting could be accelerated by fine-tuning sensor properties such as the thermodynamic stability of transient duplexes and the lengths of various spacers within the sensor. Considering the inverse relationship between the hybridization length of two oligonucleotides and the specificity of the interaction^15^, we opted to keep the query-target complementarity to 8 bp while tuning other parameters like spacer and competitor lengths to generate shorter dwell times of FRET states and more balanced distributions between FRET states.

To obtain a more balanced FRET distribution, we increased the query spacer from dT_18_ to dT_33_ (**Figure S1B**). In line with our expectations, this sensor Q_8_C_7_QS_33_CS_3_ showed a shift towards the high-FRET state (**Figure 2B, bottom left panel**). The average dwell time of the low-FRET state was 5.2 ± 0.9 s, a reduction of approximately 60% compared to the sensor Q_8_C_7_QS_18_CS_3_. In contrast, there was no significant change in the lifetime of the high-FRET state (**Figure 2C, left two panels**). A possible explanation for this observation is that increasing the length of the query spacer also increases the entropic cost of query-competitor binding, reducing the lifetime of the low-FRET state. Consistent with this hypothesis, when we reduced the query spacer to 3 nt and tested the resulting sensor Q_8_C_7_QS_3_CS_3_, a static low-FRET behavior was observed, presumably due to the minimized entropic cost of query-competitor binding (**Figure S1B & S1E).**

To further reduce dwell times, we shortened the competitor sequence from C_7_ to C_6_ and tested three different iSiMREPS sensors with the same varying query spacer lengths of dT_3_, dT_18_, and dT_33_ (**Figure 2A & S1C**) as we did with the C_7_ sensors. As with its C_7_ counterpart, the sensor Q_8_C_6_QS_3_CS_3_ did not show any high-FRET signal (**Figure S1F**). The sensor Q_8_C_6_QS_33_CS_3_ showed a significant reduction in dwell times for both high-and low-FRET states compared to its C_7_ counterpart. However, the low-FRET state dwell time was reduced substantially to 0.5 ± 0.1 s, and there was a strong dominance of the high-FRET state (**Figure 2B, top right panel**). We obtained the most promising results from the Q_8_C_6_QS_18_CS_3_ sensor (**Figure 2A**), which showed the desired near-parity between the high-and low-FRET states (**Figure 2B, middle right panel)** with shorter dwell times (relative to the sensor Q_8_C_7_QS_33_CS_3_) of 4.7 ± 0.7 s and 3.5 ± 0.4 s, respectively (**Figure 2C**). A control experiment confirmed that this sensor showed no high-FRET signal in the absence of miR-141 (**Figure 2B, bottom right panel)**. Overall, the analysis of FRET behavior and dwell time suggests that as the query spacer (QS_c_) length is increased, and the competitor (C_b_) length decreased, the query-target interaction is favored compared to query-competitor interaction. Since the Q_8_C_6_QS_18_CS_3_ sensor showed the best kinetic fingerprint, with short and well-balanced dwell times, this design was chosen for subsequent optimization and assay development for the rapid detection of miR-141.

### Monte Carlo simulations rationalize the dependence of iSiMREPS sensor kinetics on spacer length

To better understand the effect of spacer length on iSiMREPS probe kinetics, we developed a coarse-grained Monte Carlo simulation model. Herein, single-stranded (ss)DNA and double-stranded (ds)DNA strands are treated as simple worm-like chains with persistence lengths of 1.4 nm and 53 nm, respectively^24–26^. Our simulation results (**Figure S3**) show that at very short spacer lengths, the distance between the target and query strands is large (i.e., pairing is inhibited) due to conformational rigidity of the stiff anchor duplex. Increasing spacer length up to 10 nt allows the target and query strands to interact without bending the anchor duplex. Beyond 10 nt, increasing the spacer length causes the target-query distance to gradually decrease due to the query strand’s increased radius of diffusion. By contrast, the query-competitor distance decreases monotonically across all spacer lengths. The net result of these effects is that the ratio of the target-query distance and the query-competitor distance increases sharply at short spacer lengths, then gradually asymptotes to one. These results predict that increased spacer length initially yields increased binding of the target by the query probe, but yields diminishing returns as the spacer length is increased above 10 nt.

Our simulations suggest that in the limit of a very long query spacer, the distribution of the two FRET states should approach the ratio that would be predicted based purely on the Δ*G* of hybridization values of the two duplexes, which would favor the high-FRET query-target bound state. These findings (**Figure S3**) are in qualitative agreement with the experimental results for detecting miR-141 using the 6-nt competitor (**Figure 2B,C**); the QS3 sensors showed that query-target interactions were unfavorable, while the QS18 sensors showed near-parity between the two states and the QS_33_ sensors favored query-target binding.

### Optimization of an iSiMREPS sensor design for detecting ctDNA

To test for generality of the iSiMREPS approach, we next targeted a different class of nucleic acid biomarker: ctDNA. We chose an *EGFR* exon 19 deletion mutation (COSMIC ID: COSM6225; c. 2235_2249del15) [p.E746_A750delELREA]), commonly found as fragmented ctDNA in biofluids of NSCLC patients^27^. The optimized iSiMREPS sensor features the same fundamental components and architecture as the sensor design for miR-141 detection. However, it deals with the greater length and dsDNA nature of the ctDNA through two additional features. First, we added a short auxiliary probe that stably binds the extended 3^’^ end of the forward strand of the duplex mutant target DNA (**Figure 3A**) to prevent reannealing of the complementary strand once melted during sample preparation. The auxiliary probe also aims to minimize any potential secondary structure of the target strand^28^.

Second, the DNA-based architecture of iSiMREPS sensors allows us to selectively remove the capture and query probes of target-less sensors after target capture and before imaging (**Figure 1**) to reduce background. To this end, we developed a two-step process that employs a pair of ssDNA “invaders” that selectively bind and disassemble target-less iSiMREPS sensors via TMSD^29^ (**Figure 3B**). In the first step, a capture invader (CI) binds to a toehold exposed on the capture probe in the absence of target. Via TMSD, the CI disrupts the capture-anchor duplex to remove the capture probe from the surface (**Figure 3B, top panel**). This first step reveals a second toehold, which is then bound by a query invader (QI) in the second step. The QI disrupts the query-anchor duplex to remove the query probe and its fluorescent signal from the iSIMREPS sensor (**Figure 3B, top panel**). Although these invaders are designed to work on non-target-bound probes to reduce background signals significantly, the spacer on the capture probe can also act as a toehold and there is a minor probability^30^ that this can lead to removal of probes from target-bound sensors (**Figure 3B, bottom panel**).

We performed proof-of-concept studies for detecting exon 19 deletion mutant DNA using a Q_8_C_6_QS_18_CS_19_ sensor (**Figure 3A**) modeled after the best performing sensor for miR-141 detection (**Figure 2A**). For this sensor, we used a longer competitor spacer (CS_19_ versus CS_3_) to further improve parity between the FRET states. We used a query probe specific to the exon 19 deletion mutant DNA (Q_8_, *T_m_* = 23.9°C, **see Methods, and Table S2**) that was designed to maximize discrimination between the target mutant DNA (MUT) and the off-target exon 19 wild type (WT) sequence (**Figure 3A**), as predicted using NUPACK^31, 32^. For optimization of sensor designs, we used a synthetic forward strand of exon 19 deletion mutant DNA. A pair of CI and QI strands, as shown in **Figure 3A**, were designed to remove non-target-bound fluorescent probes from the surface (**Figure 3B**). However, the initial design of CI contains a single mismatch in the spacer region to prevent the use of the capture spacer as a toehold (**Figure 3A**).

To examine the performance of the Q_8_C_6_QS_18_CS_19_ sensor for detecting exon 19 deletion mutant DNA and to assess the efficacy of the invader strands, the preassembled sensor consisting of anchor strand, capture probe, and query probe was first tethered to the glass coverslip and the mutant DNA target was then introduced to bind with the sensor probes on the surface. Next the samples were (or were not) incubated with invaders and imaged with an objective-TIRF microscope. We found that invader treatment significantly reduced background signal in single-molecule intensity-time traces, resulting in a 3-fold higher signal-to noise (S/N) ratio relative to samples that were not treated with invaders (**Figure 3C & 3D**). However, inspection of single-molecule kinetic traces (**Figure 3C, top right panel**) revealed that the high-FRET state was significantly more populated than the low-FRET state, which is suboptimal because less dynamic kinetics creates less distinct fingerprints.

Exponential fitting of dwell time distributions (**Figure S4A and SI for detail**) showed average dwell times for high-and low-FRET states of 1.7 ± 0.1 s and 0.8 ± 0.2 s, respectively (**Figure 3E**). These dwell times are shorter than those measured for miR-141 detection under similar salt concentration and temperature. This change likely arose because the query-mutant DNA duplex (*T_m_* = 23.9°C) was less stable than query-miR-141 duplex (*T_m_* = 30.2°C). Moreover, the presence of the extra 3′ sequences in the exon 19 deletion target may destabilize the interaction with the query strand slightly by introducing more electrostatic repulsion from the nearby phosphates.

To modulate the dwell times of high-and low-FRET states, we designed several additional iSiMREPS sensors. Firstly, we decreased the length of competitor spacer of the sensor Q_8_C_6_QS_18_CS_19_ to CS_12_ and CS_4_ (**Figure S5A)**. We expected that decreasing the CS length would 1) increase the rate of the query–competitor interactions because of higher local effective concentrations, and 2) increase the dwell time of the low-FRET state, making it more like that of the high-FRET state. However, the results showed that varying the CS length had an insignificant effect on the dynamics of FRET transitions in iSiMREPS sensors (**Figure 3E and S5B**). This result may have arisen because the relatively long QS (dT_18_) present in this series of designs introduced substantial flexibility to all constructs, thus undercutting attempts to finely tune effective local concentrations. Secondly, we ran experiments where we increased the length of competitors of the sensor Q_8_C_6_QS_18_CS_19_ to C_7_ and C_8_ to raise the thermodynamic stability of the query–competitor interaction (see **Table S2**). Indeed, increasing the competitor length from 6-to 8-nt increased dwell times for the low-FRET states significantly (**Figure 3F**), further confirming that competitor length is one of the most important parameters in iSiMREPS sensor design.

Overall, the Q_8_C_6_QS_18_CS_d_ = _4, 12, 19_ design, where d is the number of nucleotides in CS, worked well for the exon 19 deletion mutant DNA. Given the insignificant effect of CS length on the dynamics of smFRET signals, the choice of CS within the tested range of lengths is somewhat arbitrary; the Q_8_C_6_QS_18_CS_19_ sensor was chosen for further mutant DNA assay optimization.

### Denaturant (formamide)-assisted rapid detection of miRNA and ctDNA using iSiMREPS

Having optimized iSiMREPS designs for both miRNA and mutant DNA, we next sought to further accelerate sensor kinetics to increase the speed of kinetic fingerprinting. One possible strategy for reducing dwell times is to reduce the number of base pairs in the query-target and query-competitor duplexes. While this approach would indeed decrease the dwell times, it would also have the undesired consequence of reducing the specificity of the probe-target interactions. As a simple approach that maintains specificity, we instead chose to use the denaturant formamide, which is known to destabilize nucleic acid duplexes and decrease *T_m_* by ~2.4-2.9°C /(mol L^−1^) by interfering with hydrogen-bond formation^33^. Due to the intramolecular assembly of the iSiMREPS sensor, we still expected fast association kinetics of the probes even in the presence of denaturant.

As predicted, adding formamide (10% v/v) to the imaging buffer resulted in intensity-time traces with much shorter high- and low-FRET dwell times for both miR-141 and exon 19 deletion mutant DNA (**Figure 4A and 4D, left panels**). With a standard acquisition time of 10 s per field of view and image processing (e.g., applying thresholds for FRET intensity, signal-to-noise, and lifetimes of bound and unbound states – see **Table S3 and S4**), histograms of the number of binding and dissociation events (*N_b+d_)* for both miR-141 and exon 19 deletion DNA molecules showed good separation from background with, but not without, 10% formamide (**Figure 4A and 4D, right panels**).

**Figure 4.**
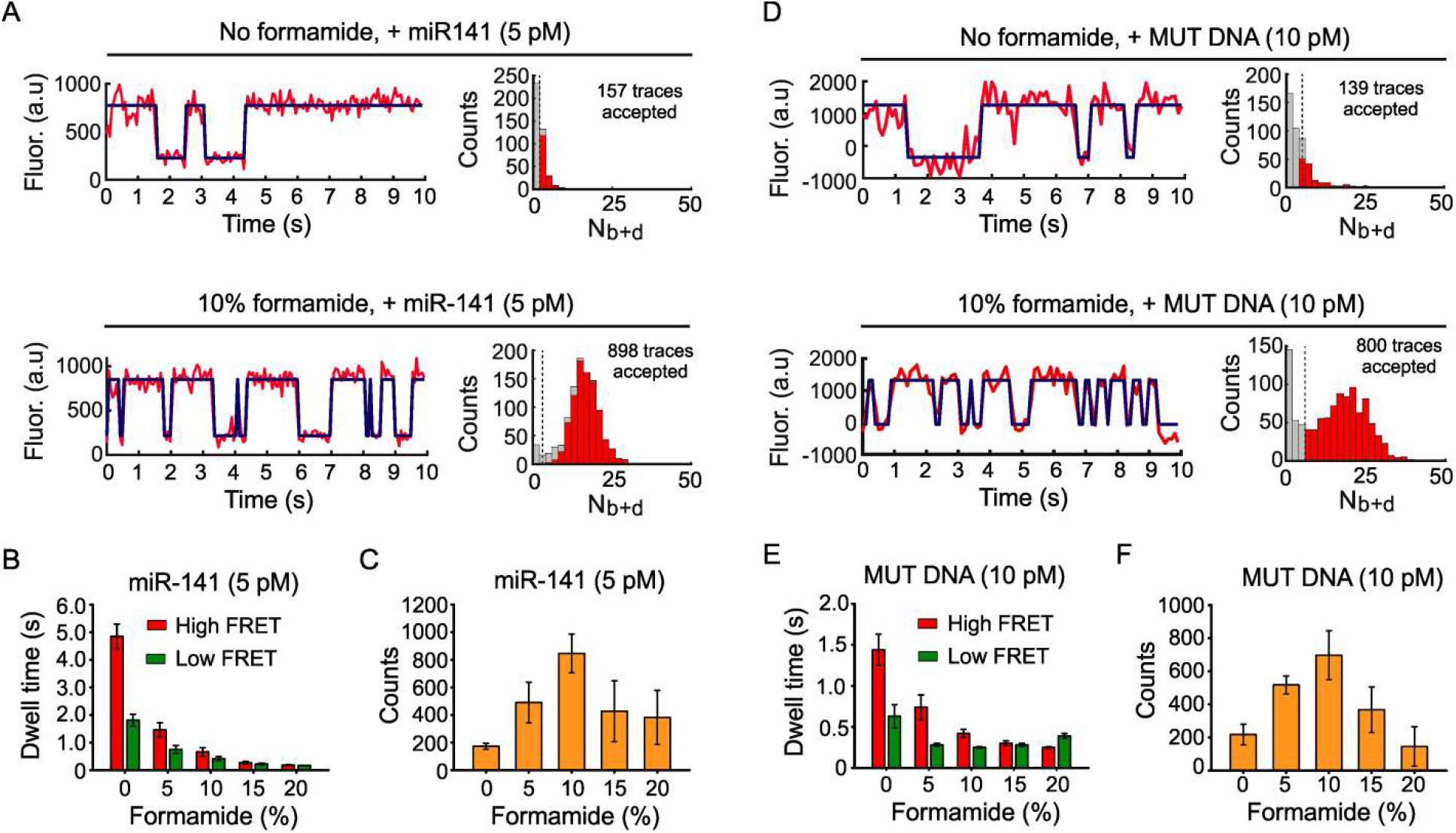
The effects of formamide on iSiMREPS sensors for rapid detection of miRNA and mutant DNA. (A) Representative single-molecule kinetic fingerprints and histograms of the number of candidate molecules per field-of-view (FOV) showing a given number of binding and dissociation events (N_b+d_) after applying thresholds for FRET intensity, signal-to-noise, and lifetimes of bound and unbound states in presence of 5 pM miR-141 in 4× PBS buffer (pH 7.4) supplemented without (top) and with 10% (v/v) formamide (bottom) at room temperature having a standard data acquisition of 10 s. The Q_8_C_6_QS_18_CS_3_— sensor as depicted in Figure 2A was used for this study and pretreated with a capture invader (5′TCCGCCATATAACACTGTCTG 3′) and query invader (5′GAGTGTCCCGCGGCCCAGGA 3′) to remove non-target-bound sensors from coverslip before imaging under an objective-TIRF microscope. (B) The average dwell times for miR-141 bound state (high-FRET) and non-bound state (low-FRET) as a function of formamide (0-20%, v/v). (C) The number of candidate miR-141 bound molecules per FOV as a function of formamide with a standard data acquisition of 10 s after applying an optimized kinetic parameter (see SI for detail, and Table S3). (D) Representative single- molecule kinetic fingerprints and N_b+d_ histograms per FOV in presence of 10 pM *EGFR* exon 19 deletion mutation in 4× PBS buffer (pH 7.4) without (top) and supplemented with 10% formamide (bottom) at room temperature having a standard data acquisition of 10 s. The Q_8_C_6_QS_18_CS_19_ sensor and invaders as depicted in Figure 3A was used for this study. (E) The average dwell times for *EGFR* exon 19 deletion mutant DNA bound state (high-FRET) and non-bound state (low-FRET) as a function of formamide (0-20%, v/v). (F) The number of candidate *EGFR* exon 19 deletion mutation bound molecules per FOV as a function of formamide with a standard data acquisition of 10 s, after applying optimized kinetic parameters (see SI for detail, and Table S4). All data are presented as mean ± s.d., where n ≥ 3 independent experiments.

Next, we varied the formamide volume fraction from 0% to 20% (v/v) to determine the shortest possible data acquisition time while retaining sensor function and high assay sensitivity. The single molecule kinetic traces showed that the dwell times for both high- and low-FRET states decreased as formamide percentage in the imaging buffer increased (**Figure S6A and S7A**). The average high- and low-FRET dwell times gradually decreased with increasing formamide in the 0-10% range for both targets, but stayed roughly constant in the 10-20% range for exon 19 deletion mutant DNA and the 15-20% range for miR-141 (**Figures S8, S9, 4B and 4E**). Specifically, shifting from 0% formamide to 10% formamide decreased average dwell times for high- and low-FRET states by factors of 7 and 4.5 respectively, for the miR-141 sensor (**Figure 4B**) and 3.5 and 2.5, respectively, for the exon 19 deletion sensor (**Figure 4E**). The slight differences between the two sensors are consistent with the fact that DNA-RNA duplexes are destabilized by formamide more significantly than DNA-DNA duplexes^34^. Plotting *Nb+d* histograms as a function of formamide showed that in a standard acquisition time of 10 s, the target bound signals separated well from background at ≥ 10% formamide and poorly or inconsistently at 0 and 5 % formamide (**Figure S6B and S7B**). The standard acquisition of ~ 10 s per field of view obtained in iSiMREPS as assisted by 10% formamide is approximately 60-times faster than intermolecular SiMREPS approaches (standard acquisition time ~ 600 s per field of view)^16, 17^.

We next determined the sensitivity of iSiMREPS sensors by calculating the number of candidate target bound molecules per field-of-view (FOV) using optimized kinetic thresholds for each formamide condition over a fixed acquisition time of 10 s. To this end, we used a Monte Carlo optimization approach to obtain a set of kinetic threshold values that minimizes acceptance of false positives in a negative control dataset while maximizing true positives in a data set with the target nucleic acid^35^ (see **Table S3 and S4)**. For miR-141 and exon 19 deletion mutant DNA, the accepted counts per FOV (a measure of assay sensitivity) increased as formamide percentage increased, reaching a maximum at 10% formamide before decreasing again at higher formamide concentrations (**Figure 4C and 4F**). The accepted counts for 0 and 5% formamide likely underrepresented the number of true molecules because many target-bound sensors could not be effectively differentiated from the background in the 10 s data acquisition period **(Figure S6B and S7B**). The lower number of counts observed in 15 and 20% formamide likely occurred for different reasons: Firstly, the reduced stability of the duplexes at these percentages suggests a higher possibility of the sensors denaturing over time; secondly, the true dwell times at higher percentages may have been too short to record with the 60-100 ms exposure times that we used in our recordings, thus reducing the observed S/N and causing a higher rejection rate of genuine target molecules (**Figure S6C and S6D, S7C and S7D**). Overall, we achieved rapid detection (~10 s per field of view), better discrimination of signals from target-bound sensor from background, and a maximization of counts per FOV for both miR-141 and exon 19 deletion mutant DNA using 10% formamide. We thus used 10% formamide during subsequent sensor optimization aimed at achieving high sensitivity and specificity.

### Use of invaders to increase the sensitivity of iSiMREPS

Our previous experiments showed that use of invaders for TMSD in the iSiMREPS assay significantly reduced background signals and improved S/N in single-molecule kinetic traces (**Figure 3**). However, we sought to better quantify the effect invaders have on the number of counts per FOV that pass kinetic filtering. Firstly, we performed a set of five experiments with the optimized sensor for exon 19 deletion DNA, each pairing the query invader shown in **Figure 3A** with one of five different capture invaders shown in **Figure 5A**. These capture invaders have different toehold and pairing region lengths. Some also contain mismatches to the spacer region of the capture probe, which are intended to mitigate undesired displacement of capture strands from target-bound sensors. We also performed a control experiment without invaders. These experiments showed that all five capture invaders increase the number of detected counts per FOV (**Figure 5B and C**) and decrease the number of false positives in a control without mutant DNA; while approximately 10 false positives per FOV were observed without invaders, there were approximately zero false positives with the invaders. However, treatment with capture invaders that contain one or more mismatches (CI_17_, CI_18_, and CI_21_) with the capture probe’s 3 nt spacer showed many more accepted traces and, surprisingly, improved S/N compared to treatment with fully complementary capture invaders (CI_20_ and CI_15_) (**Figure 5B & C, and S10**). These results suggest that fully complementary capture invaders cause unwanted removal of target-bound probes. Overall, treatment with the invaders CI_17_ and QI showed the most consistently high number of accepted traces and the best S/N, increasing the number of accepted traces ~4.5-fold compared to assays that did not use invaders (**Figure 5C & D**).

**Figure 5.**
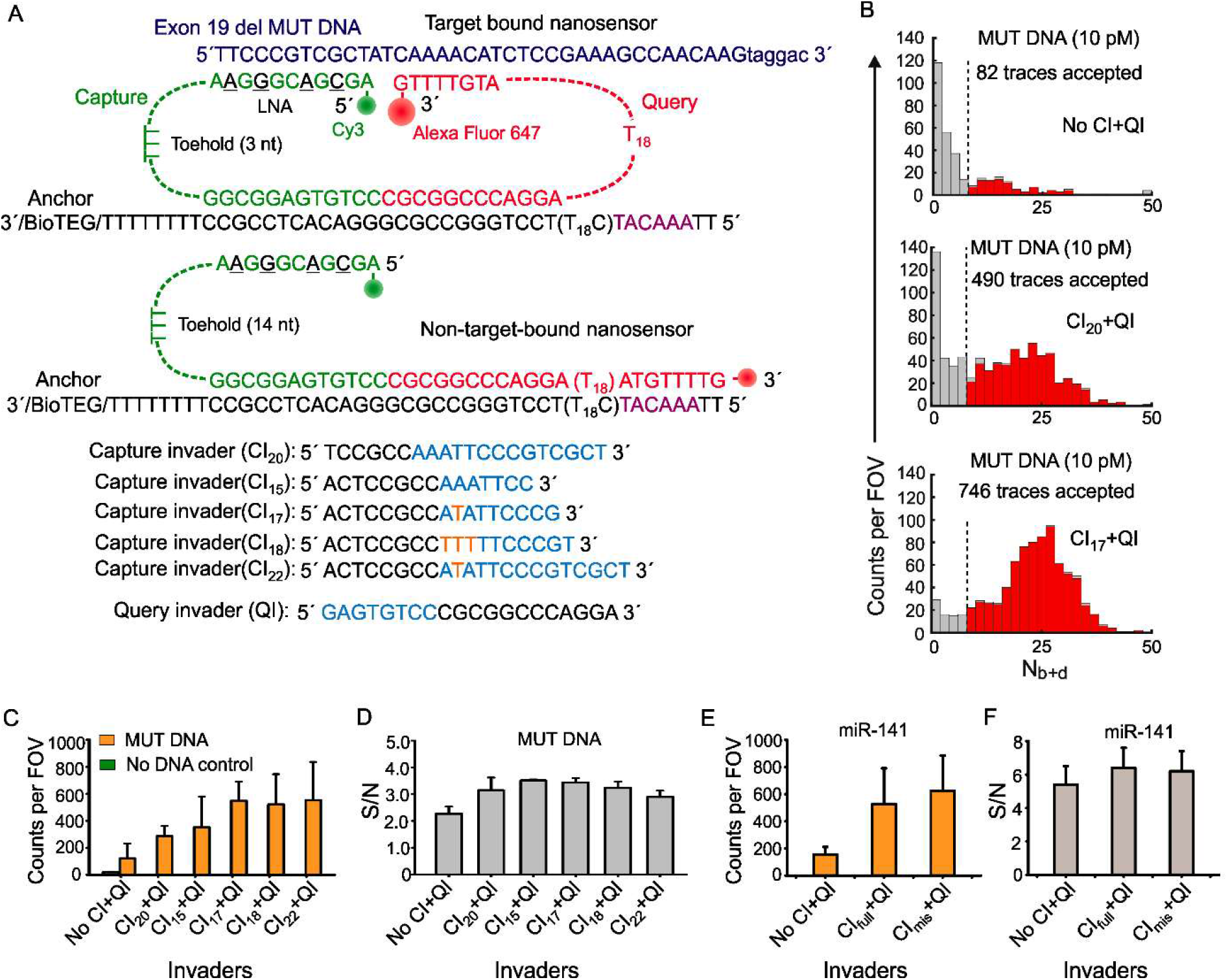
Optimization of the invaders for increased sensitivity of iSiMREPS assays for nucleic acids. (A) Schematic of target bound and non-target-bound iSiMREPS sensors, depicting the toehold available for invader binding as well as capture invaders of variable lengths. Cyan segments of the invaders are complementary to the exposed toeholds, while orange sequences represent the nucleotides that are mismatched between invaders and toeholds in the capture probe. (B) Histograms of the number of candidate molecules per field-of-view (FOV) showing a given number of binding and dissociation events (Nb+d) detected in 10 s per field-of-view (FOV), after applying thresholds for FRET intensity, signal-to-noise, and lifetimes of bound and unbound states without (top) and with (middle and bottom) the application of different invaders. (C) and (E) Number of accepted counts per FOV in the presence of *EGFR* exon 19 deletion mutant DNA and miR-141 respectively, after application of different capture invaders. (D) and (F) Signal-to-noise (S/N) ratio in the candidate target bound molecules after application of different capture invaders. All data are presented as the mean ± s.d. of n = 3 independent experiments.

This strategy was also tested and optimized for the detection of miR-141. Specifically, one experiment without invaders, one with a fully complementary CI, and one with a capture invader with one mismatch in the region complementary to the capture spacer were performed to test if miR-141 followed a similar pattern of optimization (**Figure S11**). The results indicated that using invader strands increased the accepted counts by a factor of ~3.5 (**Figure 5E**). The mismatched CI performed better than the fully complementary one and showed a modest improvement in the number of accepted traces (**Figure 5E**) and the S/N (**Figure 5F**). This can be attributed to the lower sensor concentration required for the miR-141 experiments, where the sensor was assembled with miR-141 in solution. Overall, we infer that the use of a pair of invaders to remove sensors lacking a nucleic acid target increased sensitivity by reducing background noise.

### Sensitivity and specificity of detecting *EGFR* exon 19 deletion mutation DNA and miR-141

To further improve sensitivity for the detection of exon 19 deletion mutant DNA, we next optimized iSiMREPS preparation procedures and assay conditions (e.g., sensor concentration, invaders, and target incubation time) (see **SI, and Figure S12**). Since *EGFR* exon 19 deletion mutant DNA exists in double-stranded (ds) DNA form in biofluids, the target was thermally denatured at 90°C for 3 min and cooled at room temperature in the presence of an auxiliary probe that binds stably to the forward strand of mutant DNA (see **Methods**, and **Figure 3A**). During this step, the poly-thymine oligonucleotide dT_30_ was included in high molar excess to act as a carrier. This process was modeled after a previous protocol^17^. The auxiliary probe and dT30 help to keep the capture region of the target DNA in an ssDNA form, thus permitting efficient and specific capture of target by the surface-tethered sensor. The control experiments using mutant ssDNA and dsDNA treated with the above denaturation steps showed very similar counts, validating the protocol (**Figure 6A**). The iSiMREPS assay for the *EGFR* exon 19 deletion mutant dsDNA was found to have a limit of detection (LOD) of 3.2 fM in buffer (**Figure 6B**) and a linear dynamic range spanning ~4 orders of magnitude (**Figure S13A**), which is ~1.5 times wider than a previous conventional SiMREPS assay for the same mutant DNA using diffraction-limited data analysis^17^. iSiMREPS experiments in the presence and absence of a large (10^5^ - or 10^6^ - fold) excess of wild-type DNA showed a specificity of 99.9996-99.9999% for the exon 19 deletion, permitting detection of mutant DNA at an allelic fraction of 0.001-0.0001% (**Figure 6C, S14 and Table S5**).

**Figure 3.**
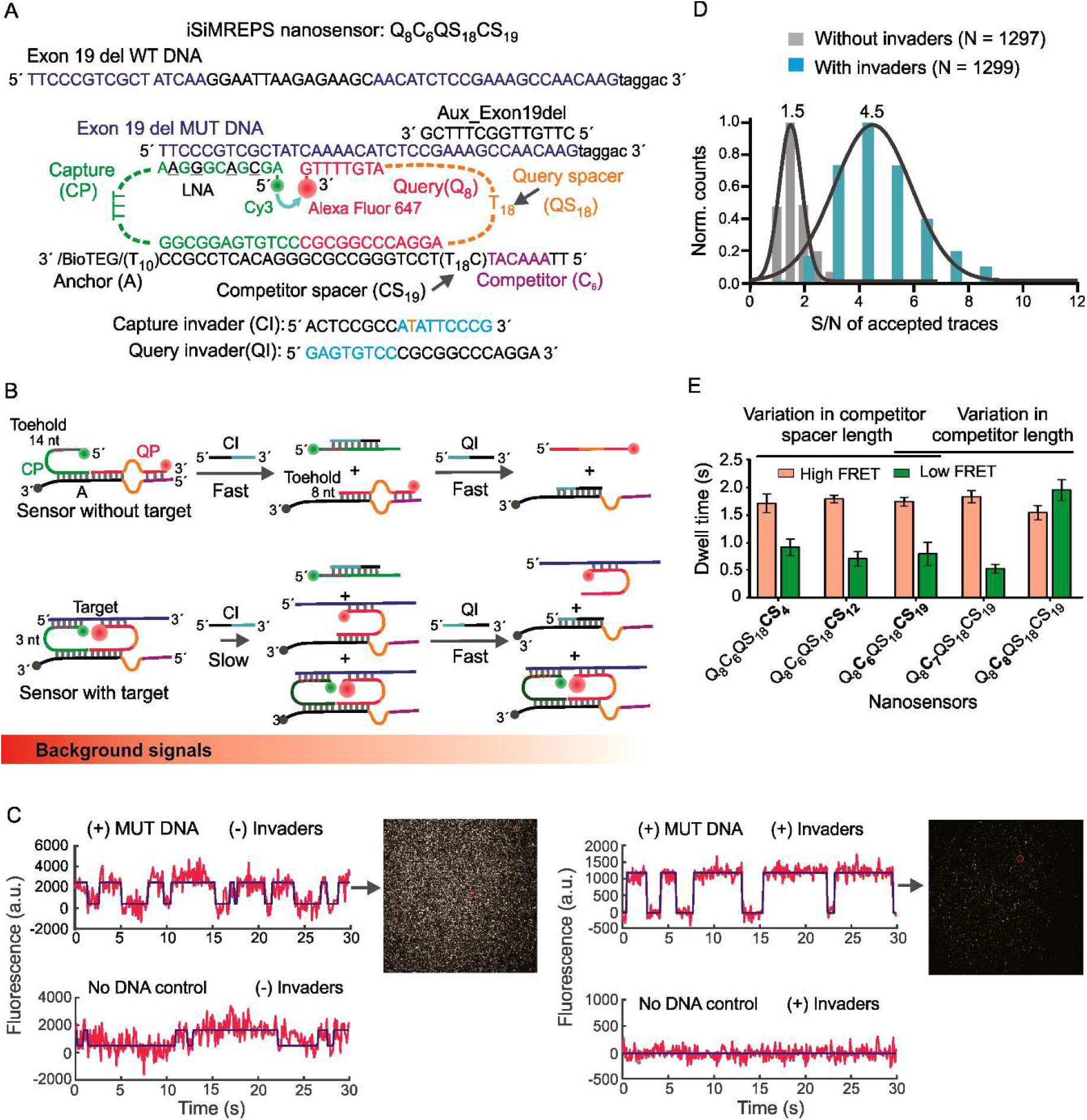
Design and optimization of iSiMREPS for detection of a ctDNA biomarker mutant DNA sequence. (A) Design of optimized smFRET-based iSiMREPS sensor for the detection of *EGFR* exon 19 deletion mutant DNA. Two invaders, which are used to remove non-target-bound fluorescent probes from the surface, are also shown. (B) Schematic depiction of the removal of non-target-bound fluorescent probes (top) using capture and query invaders, and the much slower side reaction that removes target-bound probes (bottom). Each non-target-bound sensor has an exposed 9 nt toe-hold on the capture probe that binds with capture invader (cyan) and initiates the toehold displacement cascade. A 3 nt toehold on the capture probe in target-bound sensors can also bind with capture invaders and ultimately prevent detection of a target molecule, but this reaction occurs much more slowly due to the shorter toehold. (C) Comparison of single-molecule FRET traces of iSiMREPS sensor in the presence (top) or absence (bottom) of the target sequence containing the *EGFR* exon 19 deletion. Background signals are significantly reduced with the application of invaders (right panel) compared to samples imaged without invader treatment (left panel). (D) Comparison of signal-to-noise (S/N) ratio with (cyan) and without (grey) invaders. (E) The average dwell times spent in the high-FRET (light red) and low-FRET (green) states for different iSiMREPS sensors designs. Lifetime averages were calculated using an exponential decay fitting model, where average reported was calculated from a fit single exponential distribution (if the sum squared error <0.08 and the R^2^ > 0.96) or a weighted double exponential distribution. All data are presented as mean ± s.d., with n = 3 independent experiments.

**Figure 6.**
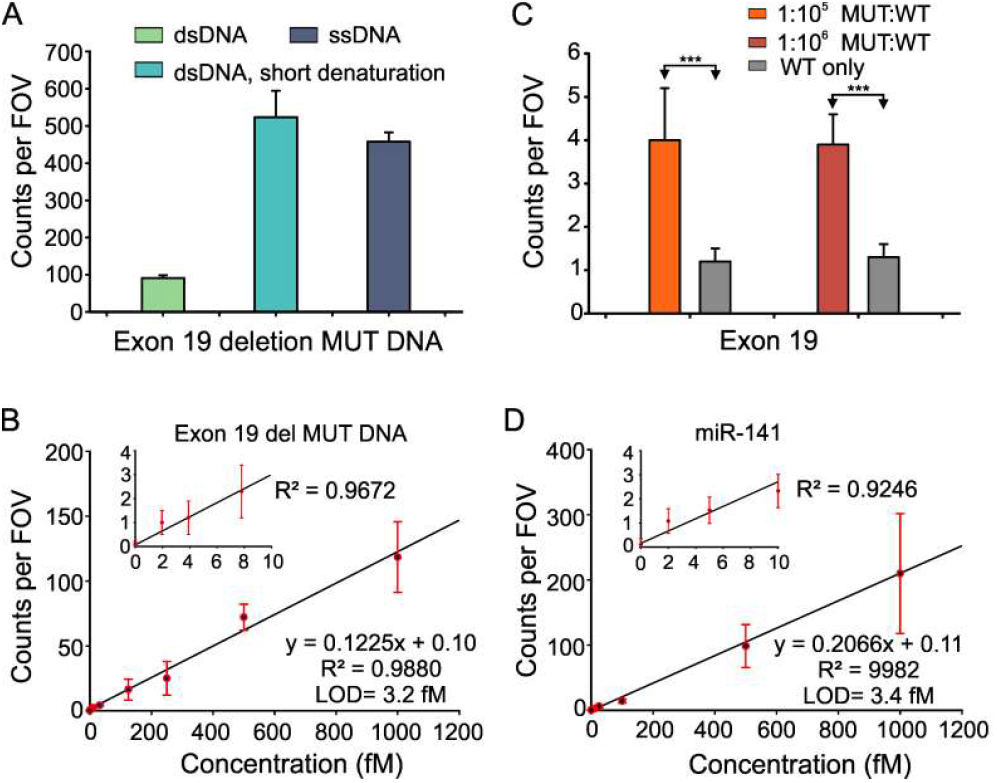
Standard curve and specificity of detecting *EGFR* exon 19 deletion mutant DNA and miR-141. (A) Effect of short thermal denaturation on the accepted counts of exon 19 deletion mutant duplex DNA. (B) Standard curve for *EGFR* exon 19 deletion mutant DNA showing a limit of detection (LOD) of 3.2 fM. Linear fits were constrained to a y-intercept of accepted counts at 0 fM, yielding an R^2^ value of 0.988. (C) Comparison of counts from low MUT allelic fraction and WT only conditions for determining specificity. Triple asterisks indicate the significant differences at 95% confidence levels as assessed using a two-tailed, unpaired t test and showed a specificity of 99.9996-99.9999% over the MUT fraction of 0.001-0.0001%. (D) Standard curve for miR-141 showing a limit of detection (LOD) of approximately 3.4 fM. Linear fits were constrained to a y-intercept of accepted counts at 0 fM, yielding R^2^ values = 0.9982. All data are presented as the mean ± s.d. of n ≥ 3 independent measurements.

For quantifying miR-141, we used assay conditions similar to these for mutant DNA with some modifications (see **Methods**). The assay for miR-141 was determined to have an LOD of 3.4 fM with a dynamic range of approximately 3.2 orders of magnitude (**Figure S13B**). This iSiMREPS-based approach showed an increase in the dynamic range of approximately 1.2-fold compared to the conventional SiMREPS approach for miR-141 detection^16^ with similar sensitivity, but in one 60^th^ of the time.

## CONCLUSIONS

We have successfully demonstrated an intramolecular smFRET-based kinetic fingerprinting technique (iSiMREPS) for direct digital counting of diverse nucleic acid biomarkers. In iSiMREPS, transient interrogation of a single target molecule using a nanoscale sensor enables rapid kinetic fingerprinting (~10 s) by dramatically increasing the local effective concentration of probes and target. The unique features of iSiMREPS permitted stable and high affinity capture of single nucleic acid molecules and high biomarker specificity. The DNA-based sensor design also allowed denaturants such as formamide to be utilized for further acceleration of probe-target kinetics; because the low-FRET state is stabilized by hybridization to a competitor sequence, formamide destabilizes both high- and low-FRET states. By contrast, in conventional intermolecular SiMREPS the duration of the unbound state would be less sensitive to denaturant. Moreover, the sensor design was amenable to a TMSD strategy – often deployed in dynamic DNA nanotechnology^29^ – that removed non-target bound sensors, thereby nearly eliminating the background noise that commonly frustrates methods for high-confidence and sensitive analysis of biomarkers.

Utilizing iSiMREPS, we have detected both miR-141 and an *EGFR* exon 19 deletion mutant DNA with a limit of detection (LOD) of ~3 fM and near-perfect specificity (>99.9999 %) within 10 s per field of view. This detection time window is approximately 60-fold faster than intermolecular SiMREPS (i.e., ~600 s per field of view) with comparable analytical performance^17^. The detection time window potentially can be further shortened with fast cameras, high laser intensities, stable fluorophores, and sensors that can withstand higher formamide concentrations and/or temperatures. iSiMREPS showed a linear dynamic range of ~3-4 orders of magnitude, which is comparable to or better than existing single-molecule techniques for nucleic acid detection^36^. While iSiMREPS has somewhat lower sensitivity than existing technologies for detecting nucleic acids like droplet digital PCR (ddPCR^10, 37^) and NGS^38^, it exhibits superior specificity, comparable or better dynamic range and faster detection of nucleic acids than ddPCR^39^, NGS^38, 40^, or Simoa^13^. Since iSiMREPS is a direct analyte counting technique, its sensitivity could be further improved by imaging many fields of view – a prospect rendered accessible by the short data acquisition time. In spite of its several advances, iSiMREPS is susceptible to the photobleaching of the fluorophores when a relatively long standard data acquisition or high laser power would be required for a particular assay. Finally, the sensitivity of SiMREPS is, as with all surface-based assays, limited by the diffusion of the target molecules to the surface. This limitation could potentially be mitigated by increasing the concentration of target molecules on the surface using a pre-concentration technique.

Overall, iSiMREPS has demonstrated the necessary analytical performance for potential applications in clinical diagnostics while still offering potential for further refinement. We anticipate that iSiMREPS can be further expanded to analyze diverse nucleic acid biomarkers beyond miRNA and mutant DNA, such as lncRNA and viral DNA or RNA, by fine-tuning the functional features of the sensors. Moreover, the intramolecular nanoscale sensor demonstrated here may be generalizable to the rapid, highly sensitive and specific analysis of diverse analytes like proteins and small molecules in a spatially addressable microarray format.

## METHODS

### Oligonucleotides and reagents

All unmodified DNA oligonucleotides used in this study were purchased from Integrated DNA Technologies (IDT, www.idtdna.com) with standard desalting purification, unless otherwise noted. Biotinylated DNA oligonucleotides were purchased from IDT with polyacrylamide gel electrophoresis (PAGE) purification. Fluorescent query probes with a 3′ Alexa Fluor 6_47_ (A6_47_) modification were purchased from IDT with high-performance liquid chromatography (HPLC) purification. Capture probes that contained locked nucleic acid (LNA) residues were purchased either from IDT with a 5^′^ C_y3_ modification and HPLC purification or from Qiagen with a 5^′^ amino modification with HPLC purification. The capture probes from Qiagen were labeled with C_y3_ monoreactive dye (GE Healthcare) and purified by ethanol precipitation and washing with 80% ethanol until the supernatant was colorless. The miR-141 with a 5^′^-phosphate modification was purchased from IDT with HPLC purification. The double-stranded exon 19 deletion mutant and wild type DNA substrates were prepared by annealing complementary single stranded oligonucleotides at 1 μM final concentration in annealing buffer (10 mM Tris-HCl, pH 8.0 at 25°C supplemented with 50 mM NaCl and 1 mM EDTA), heating at 95°C for 3 min, cooling to room temperature for 25 min, and finally keeping at 4°C for 10 min before storage at-20°C for further use. All oligonucleotides’ sequences are shown in **Table S6 and S7**.

### Preparation of slides, coverslips, and sample cells

Single-molecule fluorescence microscopy experiments were performed using either an objective-TIRF or a prism-TIRF microscope, which required different protocols for preparing slides or coverslips and sample cells as previously described^28, 41^. Objective-TIRF coverslips and imaging cells were prepared by following the protocols in three basic steps: cleaning the coverslip to remove organic residues from surface, passivating the surface with affinity tags, and preparing the sample cells by attaching cut pipette tips as described previously^17, 42^. Briefly, VWR No. 1.5, 24×50 mm coverslips (VWR, catalog no. 48393-241) were cleaned following either one of two procedures. In one cleaning procedure, the coverslips were cleaned by applying plasma for 3 min and then washed two times with acetone. In the second cleaning procedure, the coverslips were first sonicated for 10 min in acetone, then sonicated in 1M KOH for 20 min, and finally were treated with “base piranha” solution consisting of 14.3% v/v of 28-30 wt% NH_4_OH, and 14.3% v/v of 30-35 wt% H_2_O_2_ that was heated to 70-80°C before immersing the slide in it as previously described^42^. Following either cleaning procedure, coverslips were then modified to present surface amines by mounting them in a coplin jar and submerging them in a 2% v/v solution of (3-aminopropyl) triethoxysilane (APTES) (Sigma-Aldrich, catalog no. A3648-100ML) in acetone for 10 min, sonicating the jar for 1 min, incubating for another 10 min, rinsed twice with acetone, rinsed five times with water, and dried with nitrogen. Slides were then functionalized by sandwiching a 1:10 or 1:100 mixture of biotin-PEG-succinimidyl valerate and methoxy-PEG-succinimidyl valerate (Laysan Bio, Inc. catalog no. BIO-PEG-SVA-5K-100MG & MPEG-SVA-5K-1g) in 0.1M NaHCO_3_ with a final mPEG concentration of 0.25 mg/μL and a final biotin PEG concentration of 0.0025 or 0.025 mg/μL for 1:100 or 1:10 mixtures, respectively, between pairs of coverslips. To reduce nonspecific binding of nucleic acids to the surface, the remaining surface amines were quenched by sandwiching ~80 μL of 0.03 mg/μL disulfosuccinimidyltartrate (Soltec Ventures, catalog no. CL107) in 1M NaHCO_3_ between pairs of coverslips. Finally, the coverslips were dried completely under nitrogen flow and stored in the dark under air for further use for up to 3 weeks. The sample cells were prepared prior to the single-molecule experiments using 20 μL pipet tips (ART low retention, Thermo Scientific). Specifically, a razor blade was used to cut through the diameter of a pipette tip ∼2 cm from the wide end of the pipette tip and the noncut base was attached to the functionalized coverslip via epoxy (Ellsworth adhesives, hardman double, catalog no. 4001)^17^. Four pipette tips were generally attached to each coverslip in this manner. The 1:10 PEG ratio coverslips were used for objective-TIRF RNA optimization experiments and the 1:100 PEG ratio was used for all DNA and quantification experiments for both DNA and RNA. Additionally, all objective-TIRF RNA quantification experiments used plasma cleaning while all DNA experiments used piranha cleaning and RNA optimization used mostly piranha with some plasma cleaning. Both cleaning protocols showed very similar analytical performance (**Figure S15**).For prism TIRF experiments, the fluidic sample cells were constructed using two pieces of double-sided tape sandwiched between a microscope slide and glass coverslip (VWR 22×30 mm). Each microscope slide had a hole on each of two ends, which was connected to Tygon tubing for exchanging sample solutions and buffers. Prior to assembly of the sample cell, the microscope slide’s surface was cleaned using an aqueous solution of “base piranha” as described above. The microscope slides were often reused by heating the slides in warm to boiling water to loosen the glue and remove the coverslip, followed by removal of all remaining residue with a razor blade and subsequent Alconox paste and base piranha cleaning.

### Design of iSiMREPS probes

The intramolecular SiMREPS sensor design requires a stable complex of the anchor, capture, and query probes that does not dissociate from the imaging surface in iSiMREPS assay conditions. The anchor contained 12-nt segments rich in GC content (≥ 75%) to have a melting temperature (*T_m_*) of ~ 60 °C for stable hybridization with both capture and query probes. The *T_m_* between anchor and capture or query probes was estimated by IDT OligoAnalyzer (https://www.idtdna.com/calc/analyzer) using the following parameters set: target DNA concentration = 25 nM, NaCl = 600 mM, 25°C. All iSiMREPS sensors contained identical sequences in the anchor probes to stably hybridize with capture and query probes. The capture probes contained an 11- to 12-nt target-capturing sequence with 4 LNA residues (*T_m_* = ~70°C, estimated using Qiagen web application) for high affinity and kinetically stable capturing of nucleic acid targets on the surface. All query probes used a 8-nt complementary section for transient binding and dissociation with miR-141 (*T_m_* = 30.2°C) (**Table S1**) or *EGFR* exon 19 mutant DNA (*T_m_* = 23.9°C) (**Table S2**) deletion mutant DNA targets. The query probes also used a 6-7-nt complementary section for transient binding with the competitor sequences extended from anchor for miR-141 (*T_m_* = 7.5 to 18.1°C) (**Table S1**) and a 6-8-nt complementary section for *EGFR* exon 19 (*T_m_* = 0 to 23.9°C) (**Table S2**). The *T_m_* between query and target or competitor was estimated by IDT OligoAnalyzer as before, but with target RNA or DNA concentration = 1 μM. The discrimination between mutant (MUT) and wild-type (WT) DNA with a specific query probe was calculated using the web software NUPACK ^31, 32^ and utilizing the following equation 1^17^,

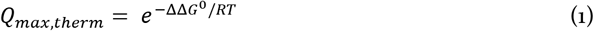

where *Q*_*max,therm*_ is the maximum theoretical discrimination, ΔΔ*G*^0^ is the difference in the Gibbs free energy of hybridization of a query with MUT and of the same query with WT DNA target. The detailed guidelines for designing SiMREPS query or fluorescent probes have been discussed elsewhere^16, 17, 28^.

### Prism-type TIRF iSiMREPS assay for detection of miR-141

To detect miR-141 using a prism-TIRF microscope, a fluidic sample cell was first passivated by injecting 150 μL of 1 mg/mL biotin-BSA (Thermo Fischer, 25mg ImmunoPure) for 10 min to coat the slide surface with biotin-BSA. The chamber was then washed out with T50 (10 mM Tris pH 8.0 at 25°C, 50 mM NaCl) and 150 μL of streptavidin at 1mg/mL concentration was flowed into the chamber, and the streptavidin was allowed to incubate for 10 min to bind with the biotin-BSA. The unbound streptavidin was then washed out with 4× PBS (Phosphate-buffered saline, pH 7.4 at 25°C). Next, 150 μL of preassembled iSiMREPS sensors bound with miR-141 were injected into the chamber for tethering onto the slide surface via biotin-streptavidin linkages. The sensors used for this step were assembled by combining the anchor, capture, and query probes as well as the miR-141 target at 1.000:1.125:1.125:1.250 ratios respectively at approximately 100 pM final concentration in 150 μL solutions in 4× PBS buffer. After combining, the sensors were heated at 70°C for 7 min in a metal bath and then cooled at room temperature for 20 min. To prolong the lifetimes of fluorophores and thus obtain more accurate measurements of the FRET signals, an imaging buffer containing an oxygen scavenger system (OSS) consisting of 1 mM Trolox, 5 mM 3,4-dihydroxybenzoate, and 50 nM protocatechuate dioxygenase in 4× PBS was injected into the chamber prior to imaging under a prism-TIRF microscope.

### Objective-type TIRF iSiMREPS assay for detection of miR-141

For miR-141 detection experiments using the objective-TIRF microscope, sample cells made of cut pipette tips were attached to a biotin-PEG and mPEG passivated glass coverslip. The sample cells were first treated with 45 μL of 0.1-0.2 mg/mL streptavidin in T50 buffer for 10-20 min. The subsequent steps for this assay followed one of two procedures. In one procedure, the anchor, capture, query probes and miR-141 target were combined at 200, 225, 250 and 5 nM final concentrations in 4× PBS buffer, heated at 70°C for 7 min in a metal bath, and then cooled at room temperature for 25 min. Unless otherwise noted, all nucleic acid samples preparation were performed in GeneMate low-adhesion 1.7 mL micro centrifuge tubes in 4× PBS. The sensor was diluted 1000-fold in 4× PBS, and 100 μL of the sensor solution was added to the cell for 45 min to tether the sensor on the surface via streptavidin-biotin affinity linkages. After removing non-surface-bound sensors and washing the cell 3 times with 4× PBS, a 100 μL solution of a pair of invader strands (see **Table S6**), each at 2 μM, was added to the cell and incubated for 20 min to remove non-target-bound fluorescent probes from the imaging surface. Next, the invader strand solution was removed, the cell was washed 3 times with 4× PBS, and 200 μL imaging buffer containing OSS in 4× PBS was added in the cell which was then imaged by TIRF microscopy. The above procedure was followed for the initial optimization of iSiMREPS assay parameters and conditions for detecting miR-141. In the second procedure, the anchor, capture, and query probes were combined in a PCR tube at 400 nM final concentrations in 4× PBS, heated at 95°C for 3min, 72°C for 7 min and 25°C for 25 min and 4°C for 10 min using a thermocycler to form a stable intramolecular complex. The sensor was then diluted to the desired concentration of 10nM and 100 μL of the diluted sensor was added in the cell and incubated for 30 min to tether the sensor to the surface. Next, 100 μL of the miR-141 solution of desired concentrations in 4× PBS was applied in the cell for 90 min for efficient capturing of miR-141 by surface tether probes. The non-target-bound probes were removed by invaders before imaging under an objective-type TIRF microscope in presence of OSS as described above. This second procedure was followed for all RNA quantification experiments.

### Objective-type TIRF iSiMREPS assay for detection of exon 19 deletion mutant DNA

The iSiMREPS assay protocols for detecting exon 19 deletion mutant DNA (COSMIC ID: COSM6225; c. 2235_2249del15 [p.E746_A750delELREA]) were developed following similar procedures as described above for miR-141 with minor modifications and a few additional steps. Briefly, the anchor, capture and query probes were preassembled in a PCR tube at 500 nM final concentrations in 4× PBS and then heated in the thermocycler the same way as described for miR-141 above. The sensors were then diluted to 10-50 nM for tethering to a PEG-biotinylated coverslip which was pretreated with 0.2-0.5 mg/mL streptavidin for 10 min in a similar way outlined above for miR-141. The *EGFR* exon 19 mutant and the wild type samples (dsDNA or ssDNA) were prepared in a 100 μL solution at their desired concentrations supplemented with 100 nM of auxiliary probe (see **Table S7**) and 2 μM of dT_30_. These solutions were heated at 90°C for 3 min in a metal block and cooled in a water bath at room temperature for 3 min and then immediately added to the cell and incubated for 90 min to bind with the sensor probes on the surface. The non-target-bound sensors were removed with the addition of 2.5 μM invader strands for 20 min. Finally, the sample was imaged in 200 μL OSS which was prepared as outlined above using objective-TIRF microscopy. Unless otherwise noted, all optimization experiments for different sensor designs and assay conditions were carried out using a synthetic forward strand of exon 19 deletion mutant DNA, while all experiments for quantifying concentration and determining sensitivity and specificity used duplex mutant DNA.

### Single-molecule fluorescence microscopy

iSiMREPS experiments were performed using either an Olympus IX-71 prism-type TIRF microscope equipped with a 60× water-immersion objective (Olympus UPLANAPO, 1.2NA) or an Olympus IX-81 objective-type TIRF microscope equipped with a 60× oil-immersion objective (APON 60XOTIRF, 1.49 NA) with CellTIRF and z-drift control modules. An ICCD (I-Pentamax, Princeton Instruments, MCP Gain 60) or sCMOS (Hamamatsu C13440-20CU) camera was used to record movies for the prism-TIRF while an EMCCD camera (IXon 897, Andor, EM gain 150) was used for the objective-TIRF. For recording smFRET signal, the Cy3-Alexa Fluor 647 fluorophore pairs were excited by light from a 532 nm laser at a power of 15–30 mW. For reliably detecting FRET signals with satisfactory signal-to-noise, an illumination intensity of ∼50 W/cm^2^ is typically used, and the TIRF angle adjusted to achieve a calculated evanescent field penetration depth of ∼70-85 nm. Two-channel images were recorded using a prism-TIRF microscope while only acceptor channel images were recorded using an objective-TIRF microscope. In prism-TIRF imaging, the signal integration time (exposure time) per frame was 100 ms, laser power was ~ 18 mW, and movies ranging from 1-15 minutes were collected to assess FRET behavior comprehensively. In objective-TIRF imaging, the exposure time per frame was 60-100 ms, and typically 200-600 movie frames were acquired per FOV.

### Processing and analysis of prism-TIRF data

The prism-TIRF movies were processed with MATLAB scripts that detected areas of higher intensity that correspond to potential molecules and used a bead mapping procedure^41^ to pair donor and acceptor signals in both channels coming from the same molecules. These scripts generated trace files that were analyzed with other scripts, where traces that showed transitions between FRET states (indicative of fingerprint generation) were selected for further analysis of their kinetics and FRET distribution. The criteria for which traces were accepted or rejected is outlined in **Table S8**. The traces, once selected, were then further processed with MATLAB scripts to obtain FRET values and time data that could be inputted into QuB (University of Buffalo software). QuB was then used to create an idealized hidden Markov model (HMM)^22^ to assign FRET states for all traces at each time. Idealized trace data from QuB was then further processed with MATLAB scripts to do two things: (1) Obtain dwell times in the low and high-FRET states and an average dwell time per state through cumulative frequency exponential fitting (see **SI, and Figures S2**), and (2) Obtain transition occupancy density plots (TODPs) which show the frequency of molecules exhibiting transitions between particular pairs of FRET states^43^. These average dwell times and TODPs were used to evaluate the sensor performance.

### Processing and analysis of objective-TIRF data

MATLAB scripts were used to identify areas of high average FRET acceptor intensity within each field of view, generate intensity-versus-time traces from these areas, and save these traces for further analysis. These traces were then analyzed using a two-state HMM^22^ algorithm to generate idealized (noise-less) intensity-versus-time traces to identify transitions between high- and low-FRET states. Thresholds of a minimum intensity of FRET transitions as well as a minimum signal-to-noise ratio (SNR) for the FRET signal were applied to each trace to distinguish genuine FRET transitions from baseline noise^22^ (**Table S3 and S4**). Those traces passing the initial intensity and SNR thresholding were subjected to kinetic analysis to extract the number of FRET transitions per trace (*N_b+d_*), the median dwell time in the high-FRET (*τ_on,median_*), and low-FRET states (*τ_off,median_*), the intensity of the low-FRET (*I_low-FRET_*) and high-FRET (*I_high-FRET_*) states, the longest individual dwell times in the high-and low-FRET states, and the coefficients of variation (CVs) of the dwell times in the high-and low-FRET states. These extracted parameters were subjected to minimum and maximum thresholding as indicated in **Tables S3 and S4** to identify target-bound sensors based on their distinct kinetic and intensity behavior and to count the number of such target-bound sensors (“accepted counts”) observed in each movie. In addition, the cumulative frequencies of the dwell times in the high-and low-FRET states were fit to a single or double exponential function (see **SI, and Figures S4, S8 and S9**) to obtain the average dwell time in each state and generate *N_b+d_* histograms for each sensor. The *N_b+d_* histograms and average dwell times were used to evaluate the sensor’s performance in terms of separation from background and capacity for rapid detection. The accepted counts were used for quantification and assessment of sensitivity.

## ASSOCIATED CONTENT

### Supporting Information

Supporting information includes Cumulative frequency exponential fitting, Statistical mechanical simulations of sensors, Optimization of iSiMREPS assay conditions, Calculation of specificity of mutant DNA. Supporting table S1-S8. Supporting figures S1-S15.

## AUTHOR INFORMATION

### Present Addresses

Alex Johnson-Buck: 333 Jackson Plz Ste 460, Ann Arbor, MI 48103

### Author Contributions

The manuscript was written through contributions of all authors. All authors have given approval to the final version of the manuscript.

### Notes

The University of Michigan has filed patent applications related to the SiMREPS technique on which M.T., A.J.B. and N.G.W. are co-inventors. M.T., A.J.B. and N.G.W. are co-founders of a startup company, a Light Sciences Inc., which seeks to commercialize this technology. A.J.-B. is an employee of aLight Sciences Inc. The remaining authors declare no competing interests.

## ACKNOWLEDGMENTS

This work was supported by National Institute of Health (NIH) grants R21 CA204560 and R33 CA229023 to N.G.W and M.T.

## For Table of Contents Only

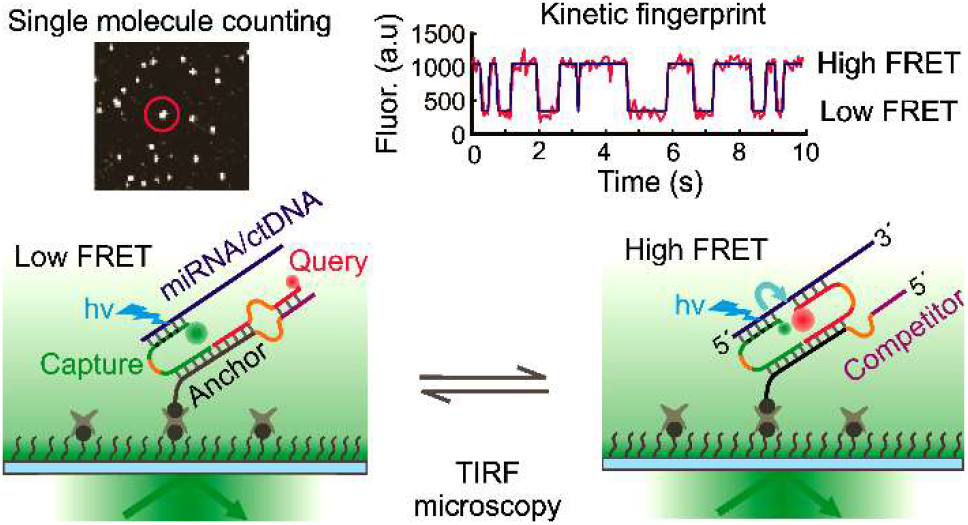

