## Supplementary Notes and Figures for "Rapid Kinetic Fingerprinting of Single Nucleic Acid Molecules by a FRET-based Dynamic Nanosensor"

AUTHOR INFORMATION

**Corresponding Author**

*

*

**Author Contributions**

The manuscript was written through contributions of all authors.

All authors have given approval to the final version of the manuscript.

Δ Kunal Khanna and Shankar Mandal contributed equally.

**Table of contents**


| Section | Contents | Page no. |
|---|---|---|
| **I** | **Supporting texts** | **3-6** |
| | 1. Exponential fitting of the cumulative frequency of dwell times | 3 |
| | 2. Statistical mechanical simulations of iSiMREPS sensors | 3-4 |
| | 3. Optimization of iSiMREPS sensor concentration, invaders, and target incubation time to increase the sensitivity of detecting *EGFR* exon 19 deletion mutant DNA. | 5 |
| | 4. Calculation of specificity for detection of *EGFR* exon 19 deletion mutant DNA. | 6 |
| **II** | **Supporting tables S1-S8** | **7-13** |
| | S1. The free energy (ΔG) and melting temperature ($T_m$) of query-target (Q-T) and query-competitor (Q-C) duplexes in different iSiMREPS sensors used for detection of miR-141. | 7 |
| | S2. The free energy (ΔG) and melting temperature ($T_m$) of query-target (Q-T) and query-competitor (Q-C) duplexes for different iSiMREPS sensors used for detection of *EGFR* exon 19 deletion mutant DNA. | 7 |
| | S3. Acquisition parameters and default kinetic filtering criteria for different iSiMREPS sensors, with and without formamide, for detecting miR-141. | 8 |
| | S4: Acquisition parameters and default kinetic filtering criteria for different iSiMREPS sensors with and without formamide for detecting *EGFR* exon 19 deletion mutant DNA. | 9 |
| | S5: Calculation of specificity for detecting *EGFR* exon 19 deletion mutant DNA. The criteria for manually selecting traces from prism-TIRF experiments. | 10 |
| | S6. The list of oligonucleotides used for detection of miR-141 | 11 |
| | S7. The list of oligonucleotides used for detection of *EGFR* exon19 deletion mutant DNA. | 12 |
| | S8: The criteria for manually selecting traces from prism-TIRF experiments. | 13 |
| **III** | **Supporting figures S1-S15** | **14-29** |
| **IV** | **Supporting references** | **30** |

### I. Supporting texts

#### 1. Exponential fitting of the cumulative frequency of dwell times

Average dwell times for a given experiment were processed using a custom MATLAB (version 2019a or later) script. The script first determined the cumulative frequency of all the dwell times for a given state using bins the size of the camera exposure time (0.06-0.1 s). This cumulative frequency was then fit to either a single exponential function (*Equation S1*) or a double exponential function (*Equation S2*):

$$y = ae^{-x/\tau} + c \tag{S1}$$

$$y = ae^{-x/\tau1} + be^{-x/\tau2} + c \tag{S2}$$

where $a$, $b$, $c$, τ, τ1 and τ2 are fit parameters. The coefficients $a$ and $b$ are used to fit the function and for the double exponential, determine the weight of each term for plotting, and obtaining average dwell times. The coefficient τ describes, for the single exponential fit, the average dwell time for a given event. The coefficients τ1 and τ2 describe, for the double exponential fit, the average dwell time for shorter- and longer-lived populations of events, respectively. The coefficient c is a constant the gives the y-intercept for the equation.

For each dataset, the cumulative frequency was first fit to the single exponential fitting function. This fit was then kept if the sum squared error < 0.05 and the $R^2 > 0.98$ for detecting miR-141 and the sum squared error < 0.08 and the $R^2 > 0.96$ for detecting *EGFR* exon 19 deletion mutant DNA, which indicated a good fit and suggested that the coefficient $t$ was an accurate average dwell time. If these conditions were not met, a double exponential function (equation S2) was used instead, and the average dwell time was calculated as $\tau = (a\tau1 + b\tau2)/(a + b)$. This equation calculated a weighted average of both populations that was reported as the average dwell time for the entire data set.

#### 2. Statistical mechanical simulations of iSiMREPS sensors

Simulations were performed using a Monte Carlo simulation method described by Becker, Rosa, and Everaers[1]. In this method, each ssDNA nucleotide (nt) or dsDNA base pair (bp) was represented as a point at fixed distance from its neighbors, $h$ (0.6 nm/nt for ssDNA[2] or 0.34 nm/bp for dsDNA[3]), and then a series of $10^7$ iterations were applied to the construct via the Metropolis-Hastings algorithm. Each iteration consisted of a pivot attempt, which entails selection of a random point in the construct, followed by a counterclockwise rotation of all downstream points (where upstream means closer to the point at which the construct is anchored to the surface) around a random axis by an angle randomly sampled from the range $\pm 50°$. The construct's post-pivot free energy, $G$, was calculated as the sum of the bending energy of all non-terminal points. The bending energy for the $i^{th}$ non-terminal point (e.g., a point that is bound to at least two additional points), $g_i$, with 3D coordinate vector $\boldsymbol{r_i}$ is:

$$g_i = -k_{s,i} \frac{(\boldsymbol{r}_i - \boldsymbol{r}_{i\leftarrow}) \cdot (\boldsymbol{r}_i - \boldsymbol{r}_{i\rightarrow})}{|\boldsymbol{r}_i - \boldsymbol{r}_{i\leftarrow}||\boldsymbol{r}_i - \boldsymbol{r}_{i\rightarrow}|} \quad (S3)$$

where $k_{s,i}$ is the point's bending spring constant, which is related to the persistence length, $L_p$ (1.4 nm for ssDNA[4] or 53 nm for dsDNA[3]), via the relation

$$L_p = \frac{-h}{\ln\left(coth(k_s) - \frac{1}{k_s}\right)} \quad (S4)$$

and $r_{i\leftarrow}$ and $r_{i\rightarrow}$ are the 3D coordinate vectors for the nearest upstream and downstream points, respectively. (Note that for single-stranded RNA in the miR-141 design, we used $L_p = 0.8\, nm$ and $h = 0.67\, nm$)[5]. Next, $G$ was calculated as $G = \sum g_i$ and the change in $G$ from the last iteration, $\Delta G$, was used to determine whether the pivot is accepted. Specifically, the pivot was accepted if $\Delta G < 0$ or, in the scenario that $\Delta G > 0$, if $\exp(-\Delta G) > R$, where $R$ is a randomly generated number sampled from the range of 0 to 1. To reflect attachment of the construct to a surface, $G$ was set to $\infty$ if any point in the construct exhibited a z-position below 0. Regardless of whether or not the pivot was accepted, the inter-strand distance was calculated at the end of each iteration as the average of the distances between the pairs of nucleotides that pair together to form the query-target duplex or the competitor-query duplex.

**3. Optimization of iSiMREPS sensor concentration, invaders, and target incubation time to increase the sensitivity of detecting *EGFR* exon 19 deletion mutant DNA.**

The iSiMREPS sensor is used to count surface-immobilized analyte molecules via a TIRF microscopy setup. Thus, the sensitivity of iSiMREPS is limited by the diffusion of target molecules to the surface, as well as the kinetics and thermodynamics of binding between the target and capture probe. The number of surface-immobilized target molecules can be enhanced by increasing the density of sensors on the surface as well incubating target solution for a longer duration. However, higher sensor density results in higher background signal, which can be addressed with higher invader concentration and/or longer invader incubation time. To increase the sensitivity of iSiMREPS, we therefore tested the performance of the sensor $Q_8C_6QS_{18}CS_{19}$ with different probe concentrations, invaders, and target incubation times.

First, the performance of the sensor was tested using 10, 25, and 50 nM sensor to detect 10 pM *EGFR* exon 19 deletion mutant DNA. The target was incubated for 90 min and pretreated with 2.5 µM invaders for 20 min to remove non-target bound sensors before imaging. The results showed that S/N values in the target bound traces decreased as sensor concentration increased (**Figure S12A**). However, the number of accepted traces per FOV was highest when 25 nM sensor was used (**Figure S12A**). This can potentially be explained as follows: with 10 nM sensor, the surface density of sensor was insufficient to efficiently capture target molecules, resulting in low counts; with 50 nM sensor, the imaging surface was saturated with target bound molecules, resulting in high background. Since 25 nM probe showed good S/N and more accepted traces per FOV, we considered this concentration for further optimization of assay conditions.

Next, we varied the incubation time of invaders (5, 10, 20, 25, and 30 min) while holding all other parameters and assay conditions equal. The number of accepted counts increased roughly linearly with the invader's incubation time and flatlined at 20 min. Therefore, 20 min was chosen as the optimized invader incubation time (**Figure S12B**). Finally, we tested the effect of target incubation time on the sensitivity of the sensor. We varied the target incubation time (30, 60, 90 and 120 min) while holding all other parameters and assay conditions equal. The results showed that the number of accepted traces increased with the target incubation time, peaked 90 min, declined at 120 min (**Figure S12C**). It is possible that at 120 min, some sensors dissociated from the surface. Therefore, a target incubation time of 90 min was chosen for further experimentation.

### 4. Calculation of specificity for detection of *EGFR* exon 19 deletion mutant DNA.

The specificity of the iSiMREPS assay for detecting *EGFR* exon 19 deletion mutant (MUT) DNA in the presence of wild-type (WT) DNA was calculated based on a previously-published protocol[6]. Briefly, to determine specificity, 500 fM MUT DNA was spiked into 50 or 500 nM WT DNA to obtain a mutant allelic fraction of 0.001 or 0.0001%, respectively. These samples were analyzed using an objective-type TIRF microscope as described in the Methods section in the main text. MUT-free samples with 50 or 500 nM WT were used as controls. The specificity was then calculated from the number of true negative (*TN*) and false positive (*FP*) counts using the following relationship.

$$Specificity = \frac{TN}{TN+FP} \tag{S5}$$

*TN* is equal to the number of WT molecules within the field of view that are not detected as MUT and *FP* is equal to the number of false positives in a WT-only experiment.

$$TN = (Number\ of\ WT\ molecules\ in\ FOV) - FP \tag{S6}$$

In an iSiMREPS assay, the number of WT molecules per field of view can be estimated by assuming that the kinetics of capture are identical for MUT and WT molecules with the equation below.

$$Number\ of\ WT\ molecules\ in\ FOV = TP \times (C_{WT} / C_{MUT}) \tag{S7}$$

$C_{WT}$ and $C_{MUT}$ are the concentrations of WT and MUT molecules, respectively, and *TP* is the number of true positives within the field of view.

$$Number\ of\ true\ positives\ (TP)\ in\ FOV = (Number\ of\ counts\ in\ MUT + WT) - (Number\ of\ counts\ in\ WT\text{-}only) \tag{S8}$$

$$TN = TP \times (C_{WT}/C_{MUT}) - FP \tag{S9}$$

By substituting equation (S9) into equation (S5), we obtain

$$\text{Specificity} = 1 - \frac{FP}{TP \times (C_{WT}/C_{MUT})} \tag{S10}$$

### II. Supporting Tables

**Table S1**: The free energy (ΔG) and melting temperature ($T_m$) of query-target (Q-T) and query-competitor (Q-C) duplexes in different iSiMREPS sensors used for detection of miR-141.

| Sensor ID | Complementary (bp) | | ΔG (kcal/mol) | | $T_m$ (°C) | |
|---|---|---|---|---|---|---|
| | Q-T | Q-C | Q-T | Q-C | Q-T | Q-C |
| $Q_8C_6QS_{18}CS_3$ | 8 | 6 | -13.56 | -9.67 | 30.2 | 7.5 |
| $Q_8C_6QS_{33}CS_3$ | 8 | 6 | -13.56 | -9.67 | 30.2 | 7.5 |
| $Q_8C_7QS_{18}CS_3$ | 8 | 7 | -13.56 | -11.62 | 30.2 | 18.1 |
| $Q_8C_7QS_{33}CS_3$ | 8 | 7 | -13.56 | -11.62 | 30.2 | 18.1 |

**Note:** ΔG and $T_m$ were calculated using IDT oligo analyzer (https://www.idtdna.com/calc/analyzer) using the complementary segments that form the duplex. All calculations were carried out at 25˚C with 1µM oligo concentrations, 600 mM $Na^+$ ions.

**Table S2**: The free energy (ΔG) and melting temperature ($T_m$) of query-target (Q-T) and query-competitor (Q-C) duplexes for different iSiMREPS sensors used for detection of *EGFR* exon 19 deletion mutant DNA.

| Sensor ID | Complementary (bp) | | ΔG (kcal/mol) | | $T_m$ (°C) | |
|---|---|---|---|---|---|---|
| | Q-T | Q-C | Q-T | Q-C | Q-T | Q-C |
| $Q_8C_6QS_{18}CS_4$ | 8 | 6 | -11.7 | -9.1 | 23.9 | 0 |
| $Q_8C_6QS_{18}CS_{12}$ | 8 | 6 | -11.7 | -9.1 | 23.9 | 0 |
| $Q_8C_6QS_{18}CS_{19}$ | 8 | 6 | -11.7 | -9.1 | 23.9 | 0 |
| $Q_8C_7QS_{18}CS_4$ | 8 | 7 | -11.7 | -10.6 | 23.9 | 11.7 |
| $Q_8C_8QS_{18}CS_4$ | 8 | 8 | -11.7 | -11.7 | 23.9 | 23.9 |

**Note:** ΔG was predicted using NUPACK[7, 8] and $T_m$ was calculated using IDT oligo analyzer (https://www.idtdna.com/calc/analyzer). The single stranded regions (spacers) flanking the complementary segments of query, target and competitor probe were considered to calculate ΔG using NUPACK[7, 8], but only complementary segments were considered to calculate $T_m$ using IDT oligo analyzer. All calculations were carried out at 25˚C with 1µM oligo concentrations, 600 mM $Na^+$ ions.

**Table S3**. Acquisition parameters and default kinetic filtering criteria for different iSiMREPS sensors, with and without formamide, for detecting miR-141.

| Parameter | Default | 0%F 10s | 0%F 30s | 5%F | 10%F | 15%F | 20%F |
|---|---|---|---|---|---|---|---|
| Frames | 1-166 | 1-166 | 1-500 | 1-166 | 1-166 | 1-166 | 1-166 |
| Exposure Time (s) | 0.06 | 0.06 | 0.06 | 0.06 | 0.06 | 0.06 | 0.06 |
| Intensity Threshold | 200 | 200 | 200 | 200 | 200 | 200 | 200 |
| Max Intensity | Inf | Inf | Inf | Inf | Inf | Inf | Inf |
| S/N Event Threshold | 2 | 2 | 2 | 2 | 2 | 2 | 2 |
| S/N Trace Threshold | 3.5 | 4.5 | 3.8 | 4.5 | 3.4 | 3.4 | 1.4 |
| Min $N_{b+d}$ | 5 | 2 | 4 | 4 | 3 | 5 | 6 |
| Max $N_{b+d}$ | Inf | Inf | Inf | Inf | Inf | Inf | Inf |
| Min Bound Median Lifetime (s) | 0.06 | 0.24 | 0.18 | 0.06 | 0.06 | 0.06 | 0.06 |
| Max Bound Median Lifetime (s) | 10 | 9.9 | 19.98 | 7.38 | 7.44 | 1.38 | 2.7 |
| Min Unbound Median Lifetime (s) | 0.06 | 0.12 | 0.18 | 0.06 | 0.06 | 0.06 | 0.06 |
| Max Unbound Median Lifetime (s) | 0.9 | 3.3 | 6.06 | 2.82 | 2.1 | 1.2 | 0.6 |
| Max Bound Event Lifetime (s) | 5 | Inf | 22.5 | 8.82 | 3.96 | 9.78 | 9.54 |
| Max Unbound Event Lifetime (s) | 4 | Inf | 15 | 5 | 4 | 4 | 2.34 |
| Max Bound CV | Inf | Inf | Inf | Inf | Inf | Inf | Inf |
| Max Unbound CV | Inf | Inf | Inf | Inf | Inf | Inf | Inf |

**Note:** All experiments other than the ones indicated specifically in this table use the settings listed under "default". The formamide variance filtering settings shown here represent data from 1 trial and were obtained using the SiMREPS kinetic parameters optimizer, which gives a starting point of filtering settings to maximize counts and minimize false positives using real and control data sets. The exact filtering settings vary from day to day for formamide experiments, as they were selected using the optimizer to gauge each condition's best possible performance.

**Table S4** Acquisition parameters and default kinetic filtering criteria for different iSiMREPS sensors with and without formamide for detecting *EGFR* exon 19 deletion mutant DNA.

| Sensors | $Q_8C_6$ $QS_{18}$ $CS_4$ | $Q_8C_6$ $QS_{18}$ $CS_{12}$ | $Q_8C_6$ $QS_{18}$ $CS_{19}$ | $Q_8C_7$ $QS_{18}$ $CS_{19}$ | $Q_8C_8$ $QS_{18}$ $CS_{19}$ | $Q_8C_6$ $QS_{18}$ $CS_{19}$ | $Q_8C_6$ $QS_{18}$ $CS_{19}$ | $Q_8C_6$ $QS_{18}C$ $S_{19}$ |
|---|---|---|---|---|---|---|---|---|
| Formamide (%) | 0 | 0 | 0 | 0 | 0 | 0 | 5-10 | 15-20 |
| Start-to-end frame | 1-200 | 1-200 | 1-200 | 1-200 | 1-200 | 1-100 | 1-100 | 1-100 |
| Exposure time per frame (s) | 0.1 | 0.1 | 0.1 | 0.1 | 0.1 | 0.1 | 0.1 | 0.1 |
| Acquisition time (s) | 20 | 20 | 20 | 20 | 20 | 10 | 10 | 10 |
| Intensity threshold per trace | 500 | 500 | 500 | 500 | 500 | 500 | 500 | 500 |
| S/N threshold per event | 1.5 | 1.5 | 1.5 | 1.5 | 1.5 | 1.5 | 1.5 | 1.5 |
| S/N threshold per trace | 1.7 | 3.7 | 2.6 | 2.9 | 4.5 | 1.5 | 1.5 | 1.5 |
| Minimum $N_{b+d}$ | 5 | 5 | 5 | 5 | 5 | 5 | 6 | 8 |
| Maximum $N_{b+d}$ | Inf | Inf | Inf | Inf | Inf | Inf | Inf | Inf |
| Minimum $\tau_{on,\ median}$ (s) | 0.1 | 0.3 | 0.1 | 0.1 | 0.1 | 0.1 | 0.1 | 0.1 |
| Maximum $\tau_{on,\ median}$ (s) | 10 | 10 | 10 | 10 | 10 | 6 | 6 | 6 |
| Minimum $\tau_{off,\ median}$ (s) | 0.1 | 0.1 | 0.1 | 0.1 | 0.1 | 0.1 | 0.1 | 0.1 |
| Maximum $\tau_{off,\ median}$ (s) | 10 | 10 | 10 | 10 | 10 | 6 | 6 | 6 |
| Minimum $\tau_{on,}$ CV | Inf | Inf | Inf | Inf | Inf | Inf | Inf | Inf |
| Maximum $\tau_{on,}$ CV | Inf | Inf | Inf | Inf | Inf | Inf | Inf | Inf |
| Maximum $\tau_{on,\ event}$ (s) | 12 | 12 | 12 | 12 | 12 | 8 | 8 | 8 |
| Maximum $\tau_{off,\ event}$ (s) | 12 | 12 | 12 | 12 | 12 | 8 | 8 | 8 |
| Maximum $I_{low\ FRET\ state}$ per trace | Inf | Inf | Inf | Inf | Inf | Inf | Inf | Inf |
| Number of intensity states | 2 | 2 | 2 | 2 | 2 | 2 | 2 | 2 |
| Ignore post photobleaching (s) | 12 | 12 | 12 | 12 | 12 | 8 | 8 | 8 |

**Note:** The default kinetic filtering criteria was determined by our newly developed machine learning based SiMREPS optimizer, which used data sets with multiple FOVs (e.g.., ≥ 10) from at least three independent experiments with and without the target as training data. For each individual experiment, the default kinetic filtering criteria were optimized slightly to minimize false positives in the negative control without rejecting true positive counts in the positive sample. The $\tau_{on}$ and $\tau_{off}$ indicate target bound (high-FRET) and non-target-bound (low-FRET) states, respectively.

**Table S5:** Calculation of specificity for detecting *EGFR* exon 19 deletion mutant DNA.

| Mutant allele (%) | MUT (fM) | WT (nM) | $C_{WT}/C_{MUT}$ | Counts ± s.d. in MUT +WT (n = 4) | Counts ± s.d. in WT-only (n = 4) | Specificity (%) = $[1 - \frac{FP}{TP \times (C_{WT}/C_{MUT})}] \times 100$ |
|---|---|---|---|---|---|---|
| 0.001 | 500 | 50 | $10^5$ | 4.0 ± 1.2 | 1.2 ± 0.3 | 99.9996 |
| 0.0001 | 500 | 500 | $10^6$ | 3.9 ± 0.7 | 1.3 ± 0.3 | 99.9999 |

**Table S6. The list of oligonucleotides used for detection of miR-141.**

| ID | Sequence: 5´-3´ | Usage |
|---|---|---|
| miR-141 | UAACACUGUCUGGUAAAGAUGG | All sensors |
| Capture_miR-141 | /5Cy3/C+A+GAC+A+GTGTTATTTGGCGGAGT GTCC | All sensors |
| Query_$Q_8QS_3$ | CGCGGCCCAGGATTTCCATCTTT /3AlexF647N/ | All sensors with $Q_8QS_3$ |
| Query_$Q_8QS_{18}$ | CGCGGCCCAGGATTTTTTTTTTTTTTTTTTCC ATCTTT /3AlexF647N/ | All sensors with $Q_8QS_{18}$ |
| Query_$Q_8QS_{33}$ | CGCGGCCCAGGATTTTTTTTTTTTTTT TTTTTTTTTTTTTTTTTTTCCATCTTT /3AlexF647N/ | All sensors with $Q_8QS_{33}$ |
| Anchor_$C_6CS_3$ | TTAGATGGTTTTCCTGGGCCGCGGGACACTC CGCCTTTTTTTT/3BioTEG/ | All sensors with $C_6CS_3$ |
| Anchor_$C_7CS_3$ | TTAAGATGGTTTTCCTGGGCCGCGGGACACT CCGCCTTTTTTTT/3BioTEG/ | All sensors with $C_7CS_3$ |
| Anchor_$C_8CS_3$ | TTTCCTGGGCCGCGGGACACTCCGCCTTTTT TTT/3BioTEG/ | All sensors with $C_8CS_3$ |
| $CI_{mis}$ | TCCGCCATATAACACTGTCTG | Removes capture probe from non-target-bound sensor. Sequence has mismatch in area that binds to capture linker. |
| $CI_{full}$ | TCCGCCAAATAACACTGTCTG | Removes capture probe from non-target-bound sensor. Sequence is fully complementary to its target on the capture probe. |
| QI | GAGTGTCCCGCGGCCCAGGA | Removes query probe from non-target-bound sensor |

**Table S7.** The list of oligonucleotides used for detection of *EGFR* exon19 deletion mutant DNA.

| ID | Sequence: 5′-3′ | Usage |
|---|---|---|
| Capture_ Exon 19 | /5AmMC6/AG+CG+ACG+GG+AATTTGGCGGA GTGTCC | All sensors |
| Query_$Q_8QS_{18}$ | CGC GGC CCA GGA TTT TTT TTT TTT TTT TTT AT GTT TTG/3AlexF647N/ | All sensors with $Q_8QS_{18}$ |
| Anchor_$C_6CS_4$ | TT AAACATC TTT TCC TGG GCC GCG GGA CAC TCC GCC TTT TTT TT/3 Bio-TEG/ | Sensor $Q_8C_6QS_{18}CS_4$ |
| Anchor_$C_6CS_{12}$ | TT AAACATC TTT TTT TTT TT TCC TGG GCC GCG GGA CAC TCC GCC TTT TTT TT/3 Bio-TEG/ | Sensor $Q_8C_6QS_{18}CS_{12}$ |
| Anchor_$C_6CS_{19}$ | TT AAACATC TTT TTT TTT TTT TTT TTT TCC TGG GCC GCG GGA CAC TCC GCC TTT TTT TT/3 Bio-TEG/ | Sensor $Q_8C_6QS_{18}CS_{19}$ |
| Anchor_$C_7CS_{19}$ | TT AAAACATC TTT TTT TTT TTT TTT TTT TCC TGG GCC GCG GGA CAC TCC GCC TTT TTT TT/3 Bio-TEG/ | Sensor $Q_8C_7QS_{18}CS_{19}$ |
| Anchor_$C_8CS_{19}$ | TT ACAAACATC TTT TTT TTT TTT TTT TTT TCC TGG GCC GCG GGA CAC TCC GCC TTT TTT TT/3 Bio-TEG/ | Sensor $Q_8C_8QS_{18}CS_{19}$ |
| *EGFR* exon 19 del MUT_ FW | TTCCCGTCGCTATCAAGACATCTCCGAAAGC CAACAAGtaggac | FW and Rev strands were annealed to prepare dsDNA. FW strand was detected |
| *EGFR* exon 19 del MUT_Rev | gtcctaCTTGTTGGCTTTCGGAGATGTCTTGATA GCGACGGGAA | |
| EGFR exon 19 WT_FW | TTCCCGTCGCTATCAAGGAATTAAGAGAAG CAACATCTCCGAAAGCCAACAAGtaggac | FW and Rev strands were annealed to prepare dsDNA. FW strand was detected |
| EGFR exon 19 WT_Rev | gtcctaCTTGTTGGCTTTCGGAGATGTTGCTTCT CTTAATTCCTTGATAGCGACGGGAA | |
| $CI_{20}$ | TCCGCCAAATTCCCGTCGCT | Removes non-target-bound capture probe |
| $CI_{15}$ | ACTCCGCCAAATTCC | Removes non-target-bound capture probe |
| $CI_{17}$ | ACTCCGCCATATTCCCG | Removes non-target-bound capture probe |
| $CI_{18}$ | ACTCCGCCTTTTTCCCGT | Removes non-target-bound capture probe |
| $CI_{22}$ | ACTCCGCCATATTCCCGTCGCT | Removes non-target-bound capture probe |
| QI | GAGTGTCCCGCGGCCCAGGA | Removes non-target-bound query probe |

**Table S8.** The criteria for manually selecting traces from prism-based TIRF experiments.

| **Criterion** | **Rationale** |
|---|---|
| Trace must have acceptor signal | Prevents traces with a bleached acceptor or no query probe from being included |
| Trace must not have multistep transitions | This convolutes the signal and makes it harder to separate genuine FRET transitions from off-target noise |
| Movies with signal that drifts into the baseline will not be accepted | Data from these movies is less trustworthy because of worsening S/N creating FRET states that can't be distinguished from noise |
| Unusually low High FRET or unusually high Low FRET values and S/N weak enough that it dips into baseline area | Traces with these features will be more susceptible to incorrect assignment of FRET states in HMM modeling |
| If there were multiple segments, the longest one was chosen and if they were of comparable lengths, the one with better S/N or clearer transitions was chosen. | This prevents the kinetic data from being too weighted or biased by a few traces with a large number of transitions. |
| If the final signal in a chosen segment is low FRET, it is only included if there is an acceptor signal after it. | This prevents signals after photobleaching of the acceptor from tainting the kinetic data |
| Traces with no distinction between baseline and signal are rejected | A static signal and an unusually intense baseline cannot be distinguished |

### IV. Supporting Figures

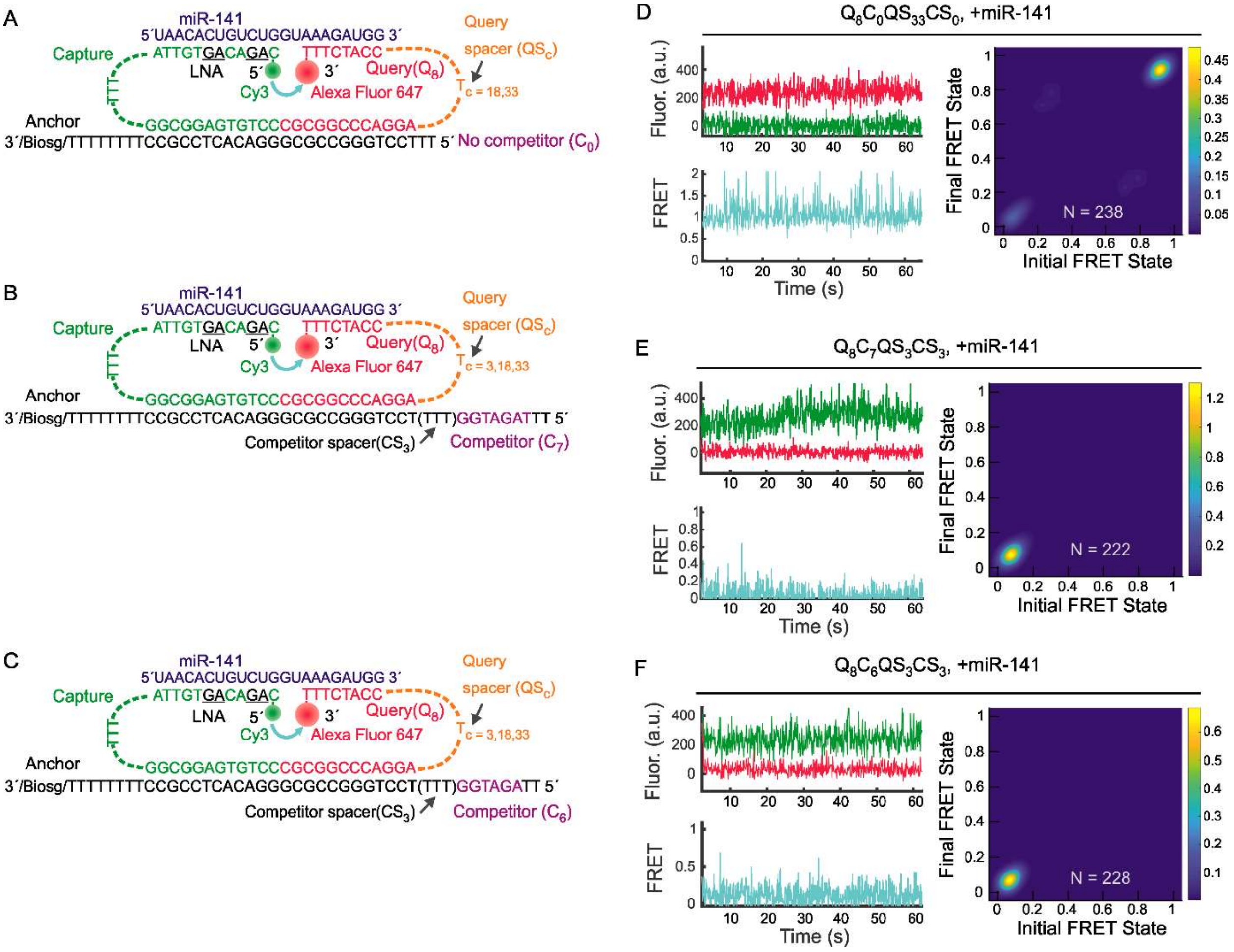


**Figure S1. Design and optimization of iSiMREPS sensors for detection of miR-141.** (A) iSiMREPS sensor designs without any competitor sequence that differ only in the length of query spacer (i.e., 18 and 33 nt). B) Sensor designs that contain a 7-nt competitor sequence that can interact with the query probe and vary in query spacer length (i.e., 3, 18 and 33 nt). (C) These iSiMREPS sensors contain a 6-nt competitor sequence and differ in the spacer lengths in the query probe. (D-F) Single-molecule kinetic traces, FRET signal, and TODP plots for the sensor $Q_8C_0QS_{18}CS_0$ (D), $Q_8C_7QS_3CS_3$ (E), and $Q_8C_6QS_3CS_3$ (F). All experiments were performed using preassembled anchor, capture, query and miR141 target at ~100 pM concentration and imaged under prism-TIRF microscopy. N represents number of molecules.

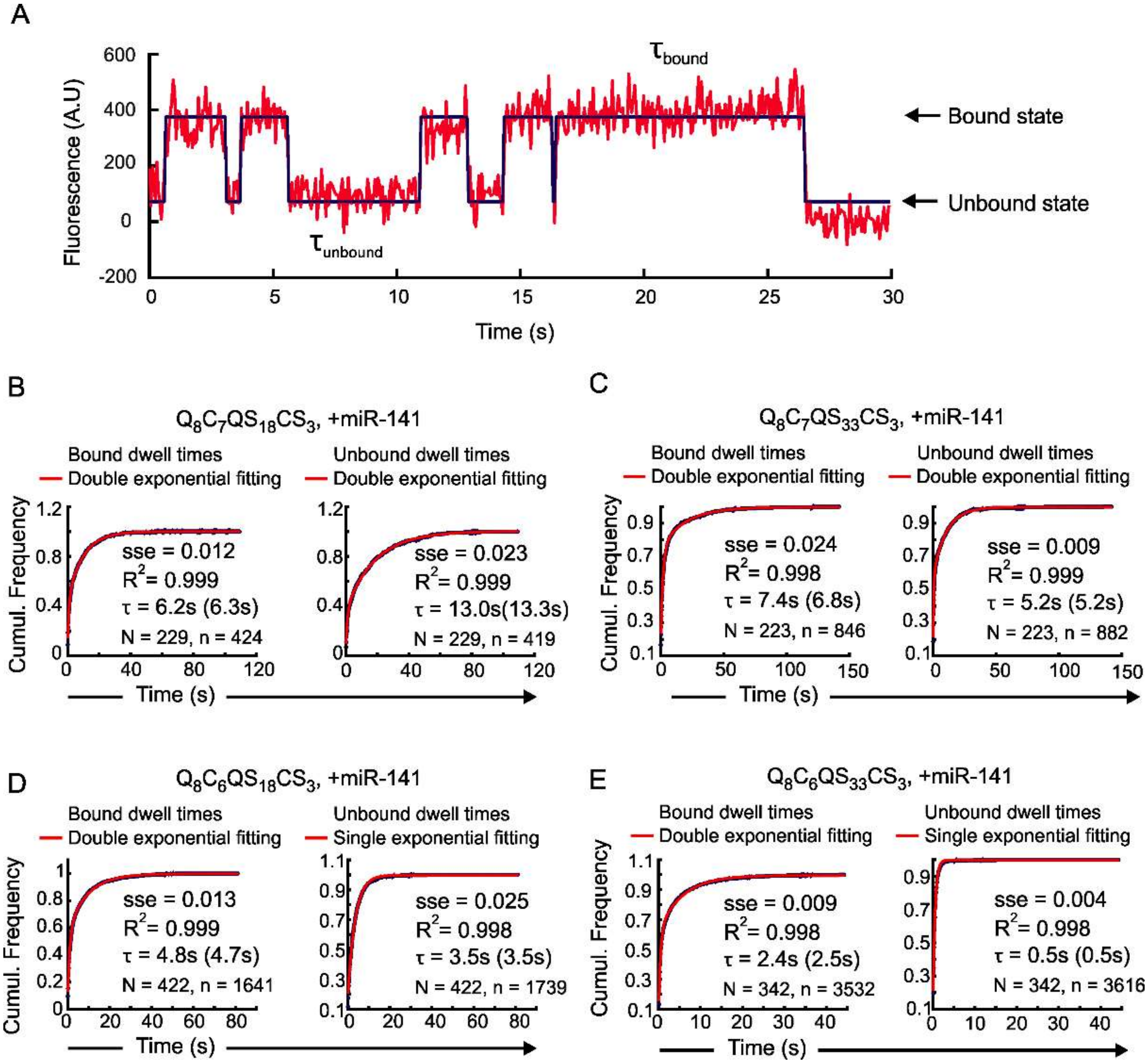


**Figure S2. Representative single molecule kinetic trace and estimation of average dwell times of FRET states for different iSiMREPS sensors for detecting miR-141.** (A) Representative intensity-time trace fitted with tan HMM to extract the dwell times of miR-141 target bound and unbound states. B-E) Exponential fitting to dwell time cumulative frequency for miR-141 target bound (high-FERT) and non-target-bound (low-FRET) states for various sensors. All experiments were performed without formamide in the imaging buffer. Single exponential fitting was chosen when sum squared error (sse) <0.05 and $R^2$ > 0.98 and double exponential fitting was used otherwise. The time listed reflects the dwell time calculated from the best-fit curve using all accepted traces, and the time in parenthesis is the reported average when the data was split into 3 populations and is the one seen in the main text. The 'N' represents number of accepted traces, and 'n' represents the total number of dwell time events used for the fitting.

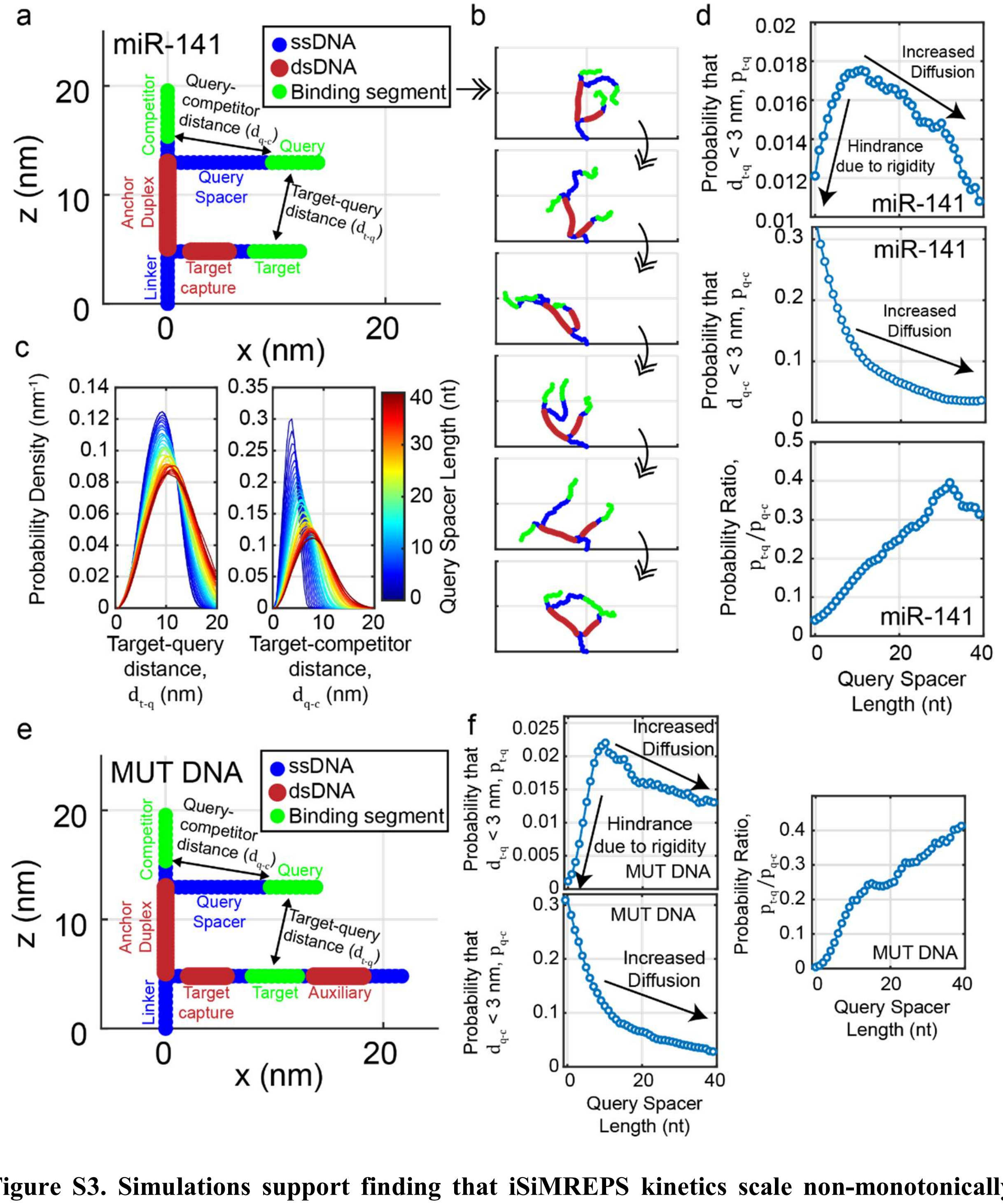


**Figure S3. Simulations support finding that iSiMREPS kinetics scale non-monotonically with query spacer length.** (A) Initialized simulated iSiMREPS construct with labels showing the three main regions of the probe (anchor, query, and target) as well as the distances between the target and query segments ($d_{t-q}$) and the query and competitor segments ($d_{q-c}$) for the miR-141 construct. All points are represented as circles with color denoted by polymer type as shown in the legend. (B) Six representative snapshots of a 2D version of the Monte Carlo simulation method, separated by at least 10,000 iterations each. (C) Probability density functions of $d_{t-q}$ for simulations with query spacers lengths (depicted by color) ranging in length from 0 nt to 39 nt. (D) Three plots are shown. The top plot shows the probability (denoted $p_{t-q}$) that $d_{t-q}$ is less than a close contact cutoff of 3 nm, as measured from the cumulative output of the Monte Carlo simulation, as a function of the query spacer length. The middle plot is similar, but for $d_{q-c}$. The bottom plot shows the ratio $p_{t-q}/p_{q-c}$. Because the activation energy for base pairing should be largely independent of spacer length and is also expected to be the rate-limiting step due to the high rate of diffusion, the strand association rate should scale linearly with $p_{t-q}$. Arrows in the top plot show that there are two roughly linear trend regimes. At query spacer lengths shorter than 9 nt, decreasing the spacer length decreases $p_{t-q}$ by what we expect is a hindrance imposed by the long, stiff anchor duplex. In this regime, it is expected that this hindrance will also increase the rate of unbinding. In contrast, $p_{q-c}$ decreases monotonically with increasing spacer length. This finding is consistent with the conformational rigidity model, as there are no dsDNA regions separating the competitor and query segments. At spacer lengths exceeding ~10 nt, increasing the spacer length mildly decreases $p_{t-q}$ due to what we expect is an increased radius of diffusion. This trend is seen for $p_{q-c}$ across the entire range of spacer lengths tested. However, while both $p_{t-q}$ and $p_{q-c}$ decrease monotonically with long spacer lengths, the ratio $p_{t-q}/p_{q-c}$ increases monotonically across the entire range, suggesting that increasing spacer length monotonically increases the preference for the target's association with the target over the competitor. These findings hold true for cutoffs that are reasonably larger or smaller than 3 nm (not shown). Notably, this simulation method is limited in that it does not account for long-range repulsive interactions between non-neighboring regions of the probe. We expect that if we did incorporate such long-range interactions, different branches of the iSiMREPS probe would be further repelled by each other, potentially steepening the correlation observed in the long-spacer length regime. (E) Initialized simulated iSiMREPS *EGFR* exon 19 deletion mutant DNA (MUT DNA) construct with labels showing the three main regions of the probe (anchor, query, and target + auxiliary complex), like that shown in A. (F) Results for a simulation of the MUT DNA design show similar trends to those shown in D.

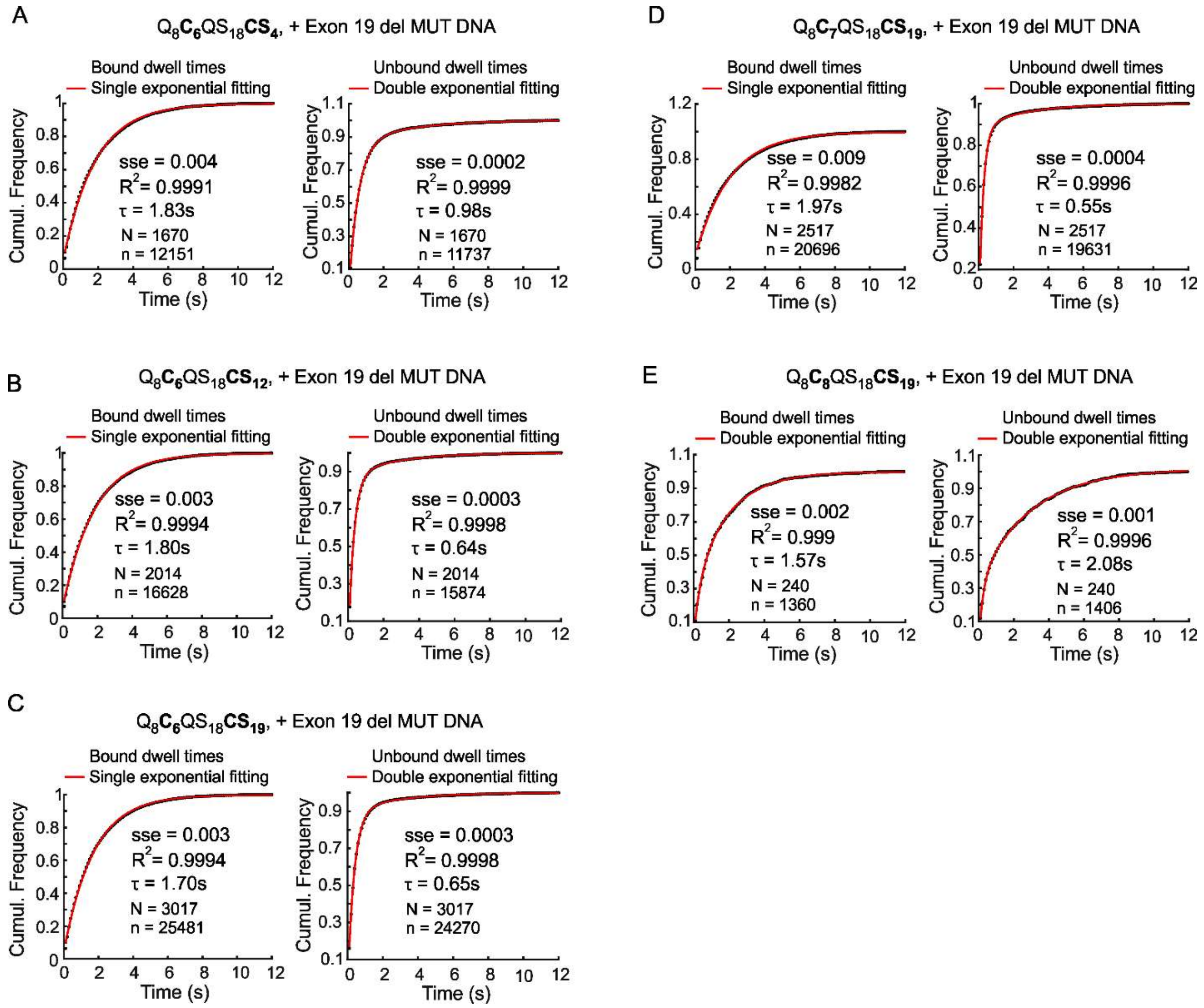


**Figure S4. Estimation of average dwell times smFRET states for different iSiMREPS sensors for detecting *EGFR* exon 19 deletion mutant DNA.** (A-E) Calculation of the average dwell time for the target bound (high-FRET) and non-target-bound (low-FRET) states for different iSiMREPS sensors for detecting exon 19 deletion mutant DNA. All experiments were performed without formamide in the imaging buffer. For all the sensors except the one with an 8-nt competitor, the target bound state dwell times were fitted with a single exponential. Single exponential fitting was chosen when sum squared error (sse) <0.08 and $R^2$ > 0.96, and double exponentials were used otherwise. All non-target-bound dwell times were fitted with a double exponential. All data is from 1 of 3 independent experiments. The 'N' represents number of accepted traces, and 'n' represents the total number of dwell time events used for the fitting.

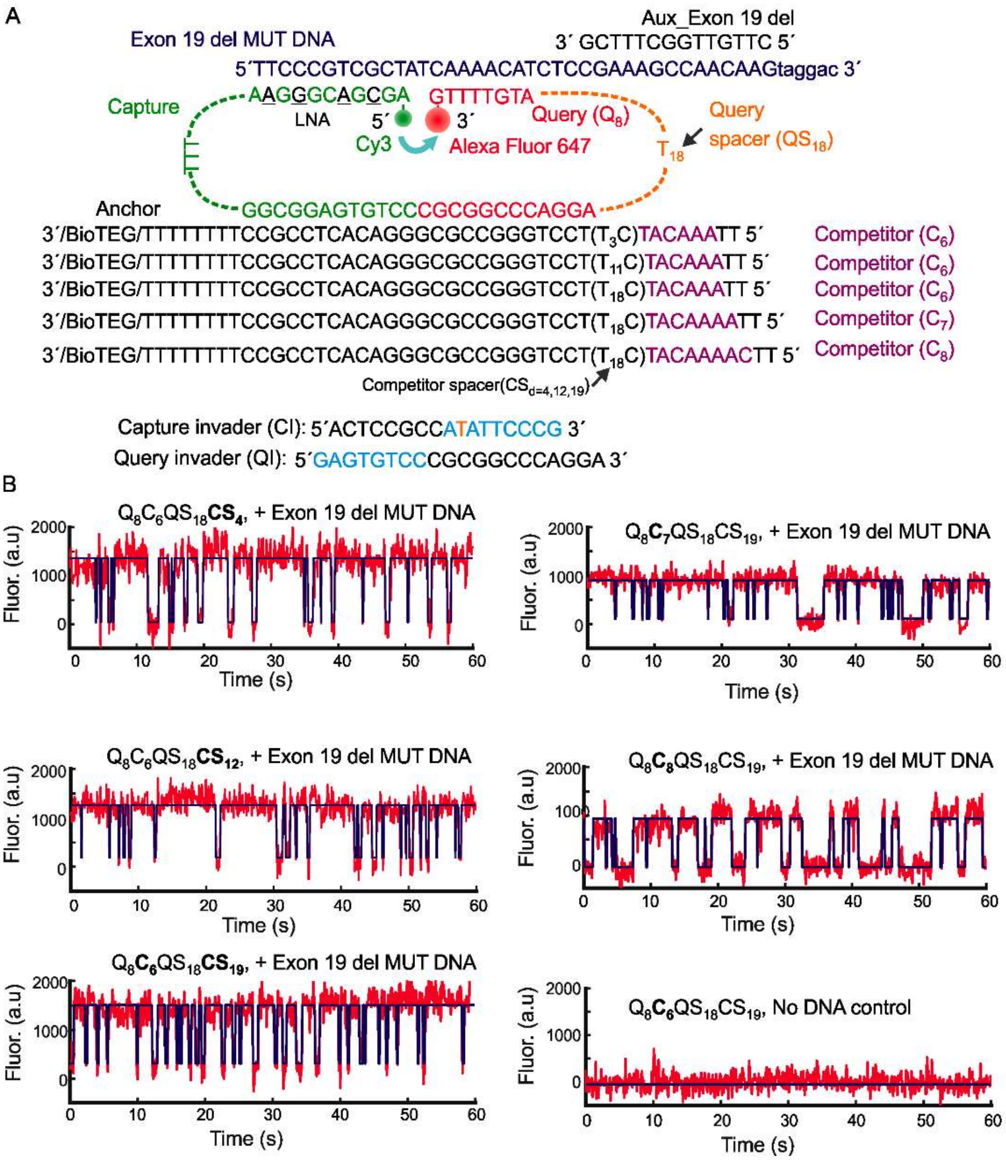


**Figure S5. Schematic of different iSiMREPS sensors and representative single molecule kinetic traces in the presence of *EGFR* exon 19 deletion mutant DNA.** (A) Designs of iSiMREPS sensors for detecting *EGFR* exon 19 deletion mutant DNA with various competitor spacer (CS) and competitor (C) lengths. (B) Representative single-molecule kinetic traces (red) for different iSiMREPS sensors with or without exon 19 deletion mutant DNA with an idealized hidden Markov model (HMM) fit (blue). All experiments were performed using 10 nM preassembled sensors consisting of anchor, capture and query probes, and 10 pM exon 19 deletion mutant DNA forward strand. Imaging was done in 4x PBS (pH 7.4) at room temperature under an objective-type-TIRF microscope. The donor fluorophore (Cy3) was excited at 532 nm and the acceptor fluorescence (Alexa Fluor 647) was recorded as FRET signal.

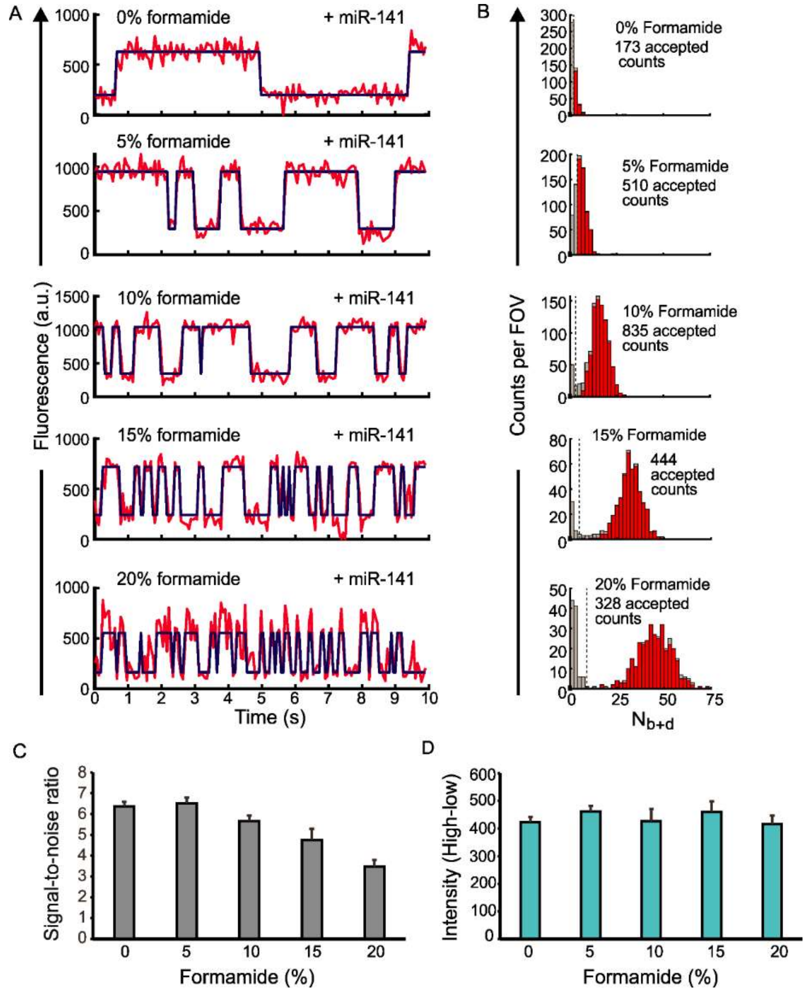


**Figure S6. Effects of formamide on the iSiMREPS sensor for detecting miR-141.** (A) Representative traces for the $Q_8C_6QS_{18}CS_3$ miR-141 sensor at 0, 5, 10, 15, and 20% v/v formamide. The signal is in red while the idealized trace obtained from hidden Markov model (HMM) fitting is in blue. (B) Histograms from 1 of 3 independent experiments for each formamide condition that show the distribution of $N_{b+d}$ among the accepted traces. These histograms reflect the distribution after application of filters for parameters such as signal-to-noise, intensity, and min and max average lifetimes. The red bars represent traces accepted while the grey bars represent traces rejected. (C) The average intensity difference between the high and low FRET states in the idealized hidden Markov model for each formamide condition. (D) The average signal-to-noise for a trace for each formamide condition. For all experiments shown, sensors were assembled at 200 nM in the presence of 5 nM miR-141. The pre-assembled sensors were then diluted it 1,000-fold and added to the surface. Imaging was performed in 4× PBS at pH 7.4. All data are presented as mean ± s.d. of 3 independent experiments.

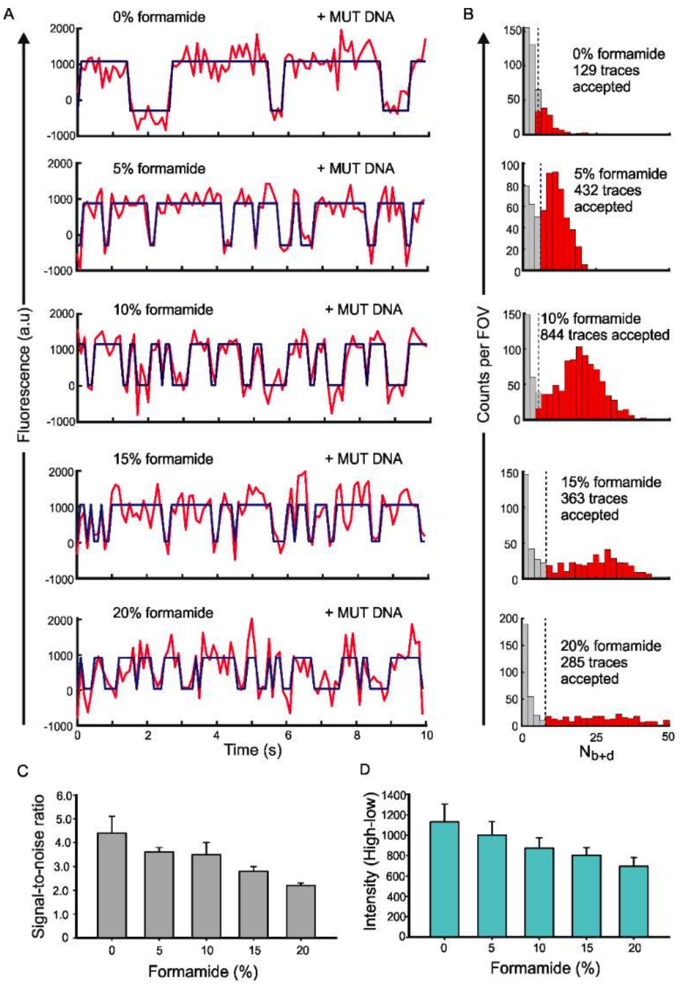


**Figure S7. Effects of formamide on the iSiMREPS sensor for detecting *EGFR* exon 19 deletion mutant DNA. (**A) Representative single-molecule kinetic traces (red) with an idealized hidden Markov model (HMM) fit (blue) of the $Q_8C_6QS_{18}CS_{19}$ sensor for detecting exon 19 deletion mutant DNA at different formamide conditions. (B) Histograms of the number of candidate molecules per field-of-view (FOV) showing a given number of binding and dissociation events ($N_{b+d}$) after applying thresholds for FRET intensity, signal-to-noise, and dwell times of target-bound and non-target-bound states for each formamide condition. Red bars represent accepted traces while grey bars represent rejected traces. (C, D) The average signal-to-noise ratio (C), and difference in intensity of high- and low-FRET states (D) of the accepted traces for each formamide condition. All experiments were performed using 10 nM preassembled sensor consisting of anchor strand, capture and query probes, and 10 pM forward strands of exon 19 deletion mutant DNA. Imaging was performed in 4x PBS (pH 7.4) at ambient room temperature under an objective-TIRF microscope. All data are presented as mean ± s.d. of 3 independent experiments.

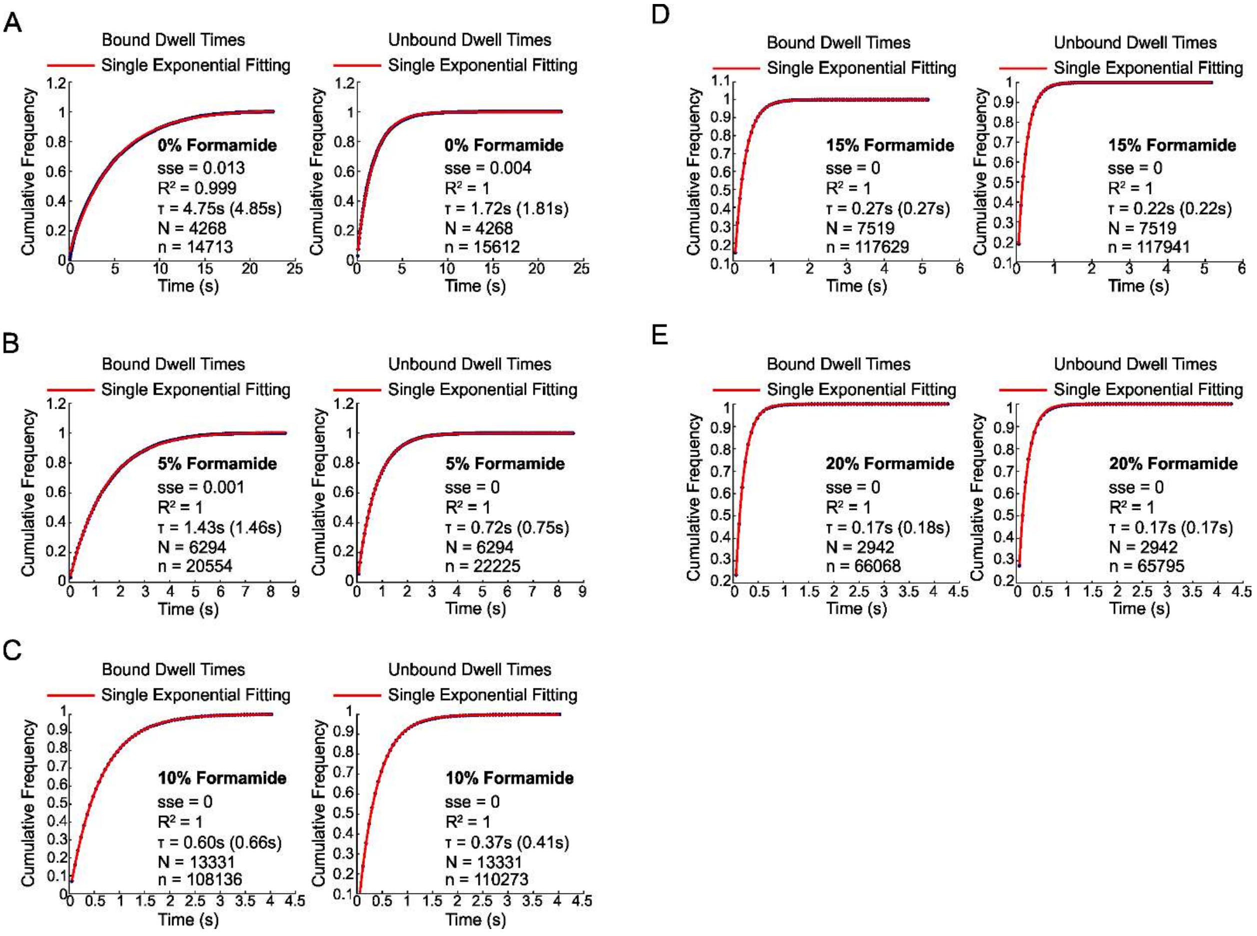


**Figure S8. Estimation of average dwell times of FRET states with the variation of formamide concentration using optimized iSiMREPS sensor for detecting miR-141.** (A-E) Calculation of the average dwell time of high and low FRET states for miR-141 for each formamide condition, obtained by fitting an exponential decay function to the cumulative frequency. Single exponential fits were used for all experiments depicted here as all had a sum squared error (sse) <0.05 and $R^2$ > 0.98. These curves represent the dwell times obtained in 1 of the 3 independent experiments conducted for each condition and the time in parentheses is the average time obtained from these independent experiments. The 'N' represents number of accepted traces, and 'n' represents the total number of apparent dwell time events in the accepted traces that used for the fitting. Experiments without formamide used a 30 s window to ensure accurate dwell times were obtained while a 10 s window was sufficient for all other conditions. The N value listed is the number of bound and unbound events in the accepted traces that contributed to the fitting.

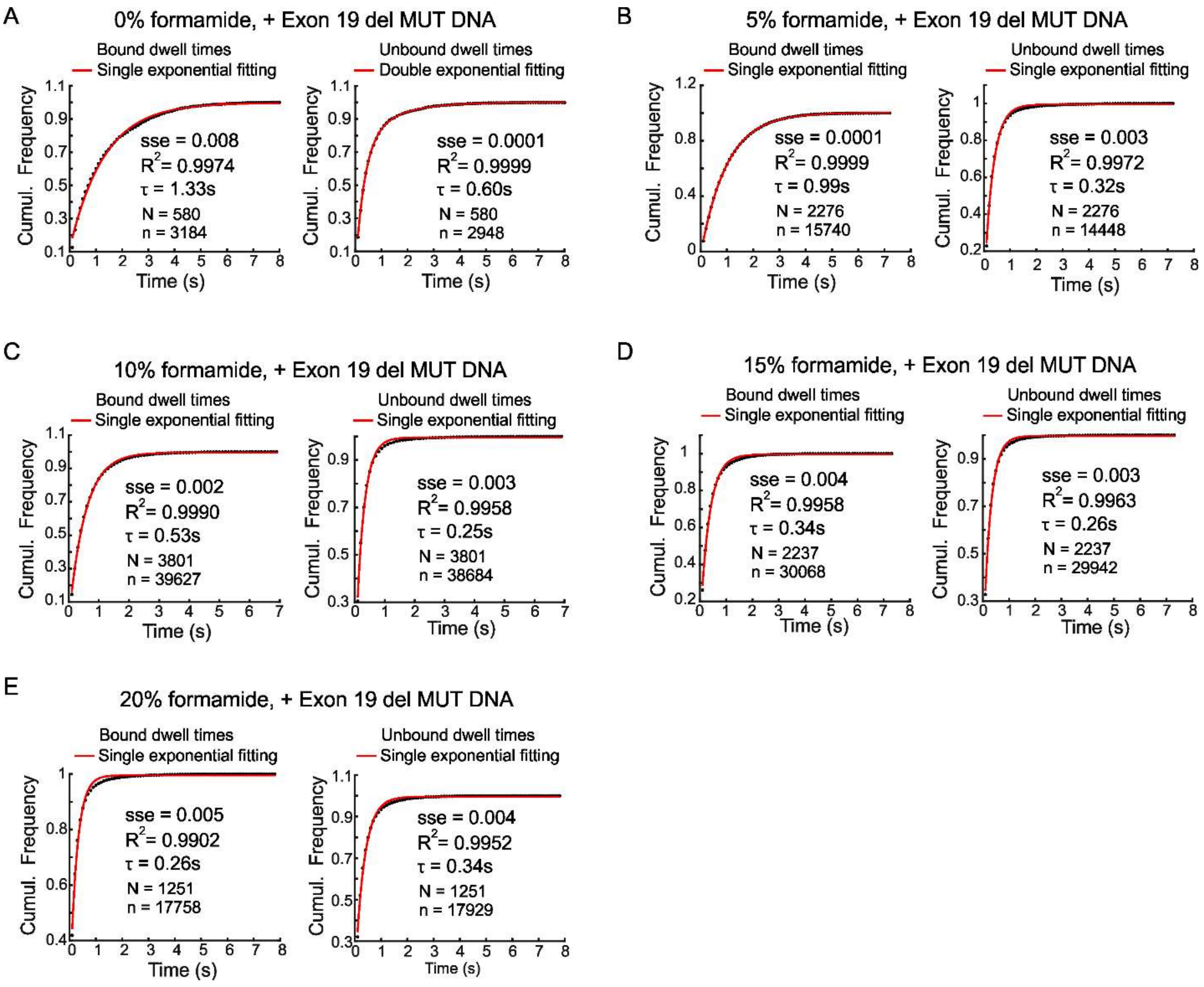


**Figure S9. Estimation of average dwell times of FRET states with the variation of formamide concentration using optimized iSiMREPS sensor for detecting *EGFR* exon 19 deletion mutant DNA. (**A-E) Calculation of the average dwell time for the target bound (high-FRET) and non-target-bound (low-FRET) states of the sensor $Q_8C_6QS_{18}CS_{19}$ for detecting exon 19 deletion mutant DNA at different formamide concentrations (0-20% v/v) by fitting an exponential decay function to the cumulative frequency. Both target bound and non-target-bound dwell times for all conditions except 0% formamide were fitted with a single exponential decay function. Single exponential fitting was chosen when sum squared error (sse) <0.08 and $R^2 > 0.96$ and double exponential was used otherwise. All data are from 1 of 3 independent experiments. The 'N' represents number of accepted traces, and 'n' represents the total number of apparent dwell time events in the accepted traces that used for the fitting.

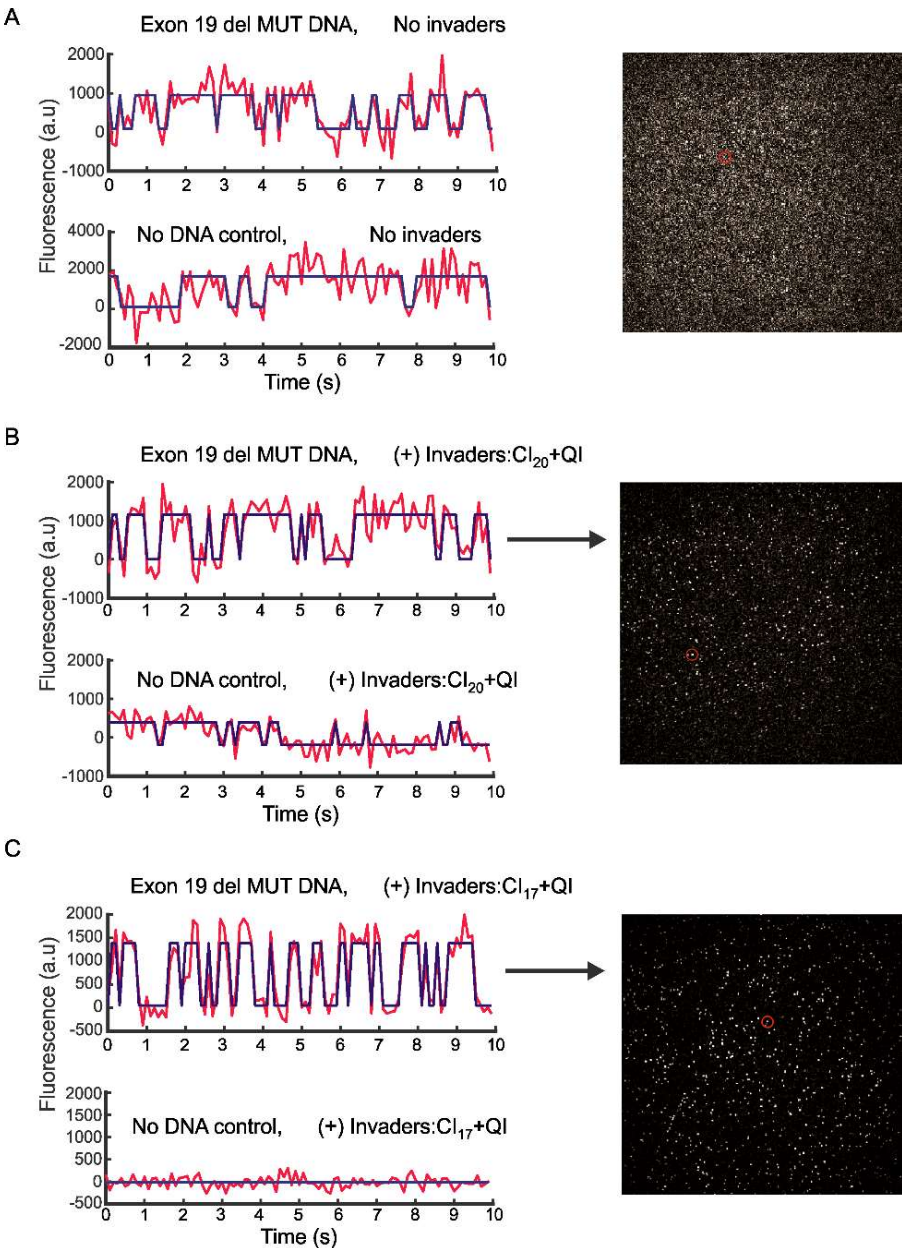


**Figure S10. Effect of different invaders on the background signals to detect *EGFR* exon 19 deletion mutant DNA.** (A-C) Representative single-molecule kinetic traces and images of a FOV without invaders (A), with invaders $CI_{20}$+QI (B), and with invaders $CI_{17}$+QI (C) in the presence and absence of exon 19 deletion mutant DNA target (see Figure 5A for invaders sequences). All experiments were performed using 0.2 mg/mL streptavidin (incubation: 10 min), 10 nM sensor (incubation: 30 min), 10 pM forward strands of exon 19 deletion mutant DNA (incubation: 90 min), 1 μM invaders (incubation: 20 min). Objective-TIRF imaging was performed in the presence of 10 % v/v formamide in the imaging buffer.

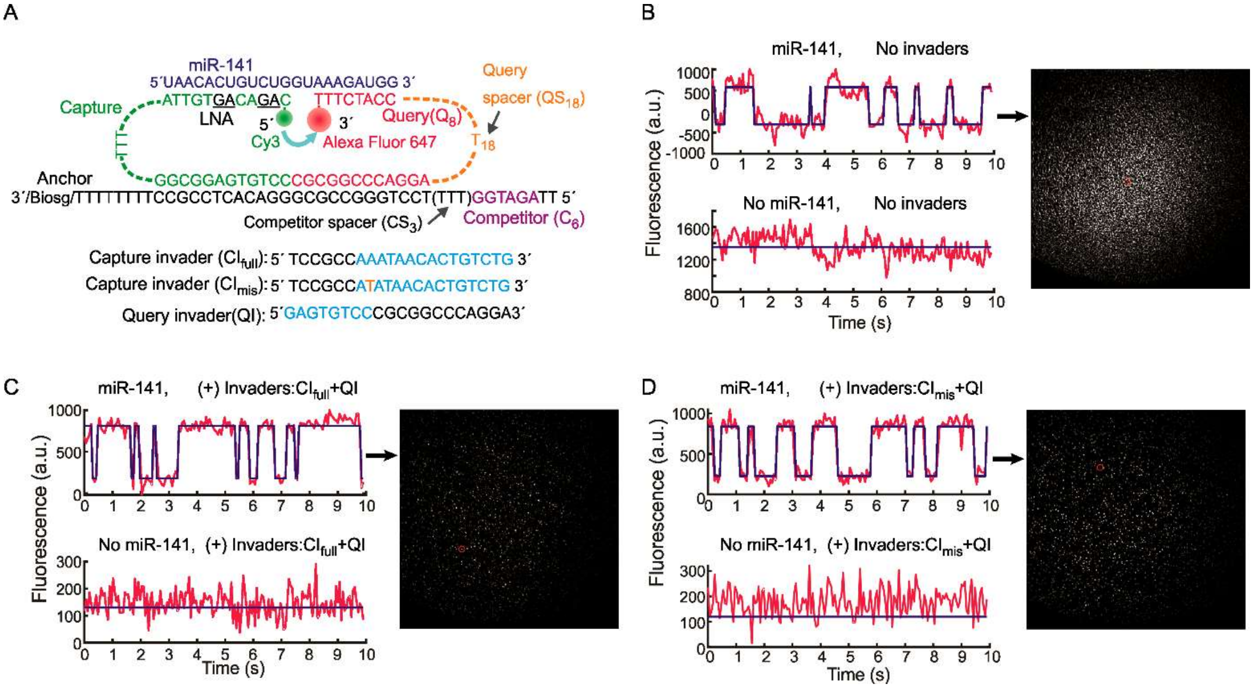


**Figure S11. Schematic of the design of iSiMREPS sensor for detecting miR-141 and representative single molecule kinetic traces in the presence and absence of different invaders**. (A) Design of the optimized miR-141 sensor and different invaders tested. (B-D) Representative single-molecule kinetic traces and images of a FOV without invaders (B), with invaders $CI_{full}$+QI (C), and with invaders $CI_{mis}$+QI (D) in the presence and absence of miR-141. Overall application of invaders improved the background signals as well the signal-to-noise ratio of single molecule traces compared to without invaders application. For all experiments shown, sensors were assembled at 200 nM in the presence of 5 nM miR-141. The pre-assembled sensors were then diluted it 1,000-fold and added to the surface. Imaging was performed in 4× PBS at pH 7.4.

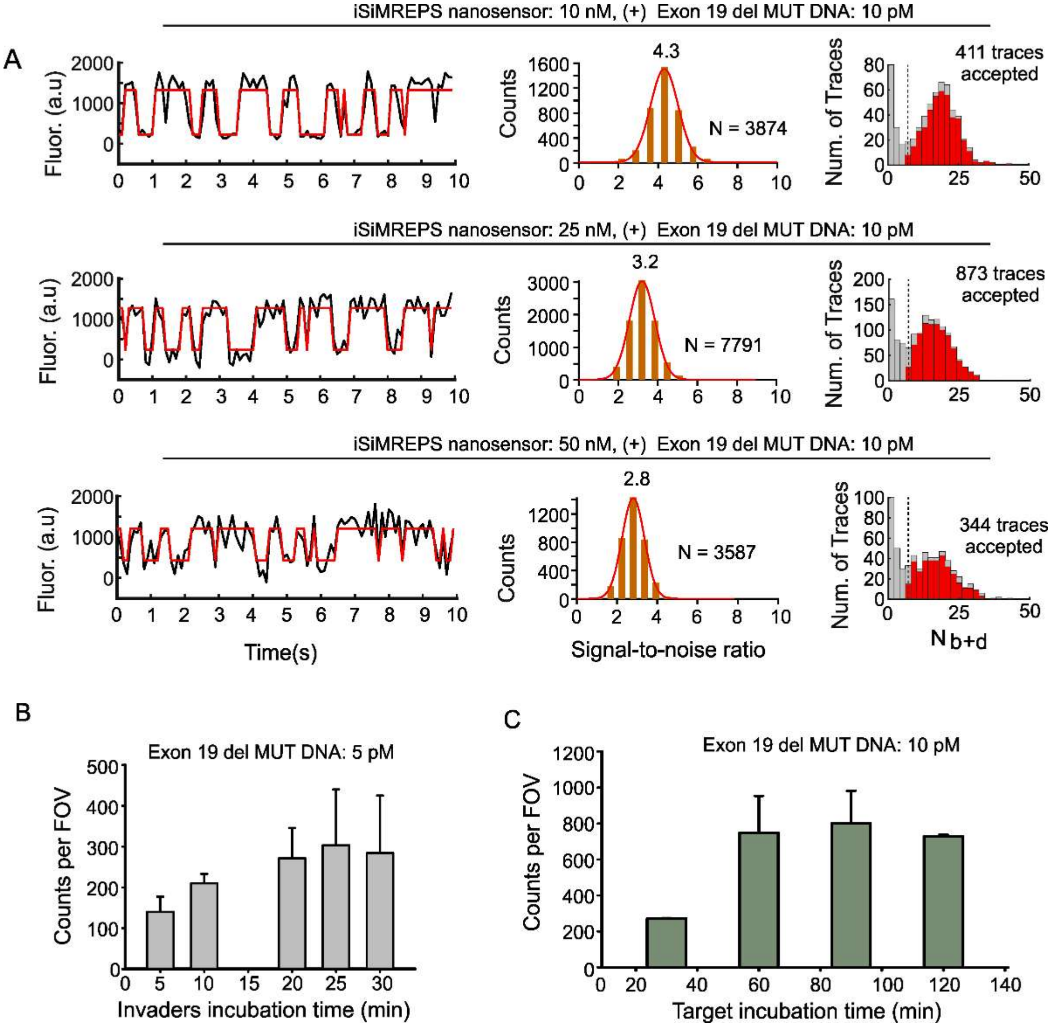


**Figure S12. Optimization of iSiMREPS assay conditions to enhance sensitivity for detection of *EGFR* exon 19 deletion mutant DNA.** (A) Effect of sensor concentration on signal-to-noise ratio (S/N) and the number of accepted traces. The experiment was performed using 10, 25, and 50 nM sensor (incubation: 30 min), 10 pM forward strands of exon 19 deletion mutant DNA (incubation: 90 min), and 2.5 µM invaders (incubation: 20 min). (B) Effect of invaders incubation times on accepted traces. The experiment was performed using 25 nM sensor (incubation: 30 min), 5 pM forward strands of exon 19 deletion mutant DNA (incubation: 90 min), and 2.5 µM invaders (incubation: 5, 10, 20, 25, 30 min). (C) Effect of target incubation times on accepted counts. This experiment was performed using 25 nM sensor (incubation: 30 min), 10 pM forward strands of exon 19 deletion mutant DNA (incubation: 30, 60, 90, and 120 min), and 2.5 µM invaders (incubation: 25 min). All experiments were performed using the sensor $Q_8C_6QS_{18}CS_{19}$. All data are presented as mean ± s.d. of 2 independent experiments.

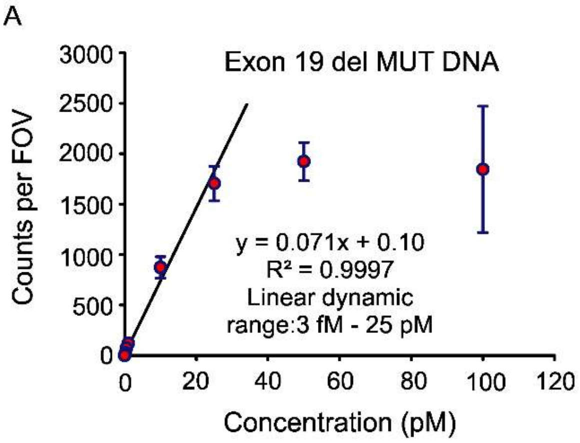


**Figure S13. Standard curves for miR-141 and *EFGR* exon 19 deletion mutant DNA.** (A) Standard curves showing the linear dynamic range for detection of *EGFR* exon 19 deletion mutant dsDNA. The experiments were performed using a glass coverslip passivated with biotin-PEG: m-PEG at a 1:100 ratio, 0.5 mg/mL streptavidin incubation for 10 min, 25 nM sensor incubation for 30 min, 1.96 fM to 100 pM exon 19 deletion mutant dsDNA incubation for 90 min, 2.5 µM invaders ($CI_{17}$ + QI) incubation for 20 min, and 10% v/v formamide. All data are presented as mean ± s.d., where n ≥ 3 independent experiments. iSiMREPS showed a linear dynamic range of approximately 3 fM - 25 pM for detecting exon 19 deletion mutant DNA which is approximately 3.9 orders of magnitude. (B) Standard curves showing linear dynamic range for detection of miR-141. The experiments were performed using a glass coverslip passivated with biotin-PEG: m-PEG at a ratio of 1:100, then incubated with 0.2 mg/mL streptavidin for 10 min, 10 nM sensor for 30 min, 2 fM to 50 pM miR-141 for 90 min, and 2 µM invaders ($CI_{mis}$ + QI) for 20 min. All imaging was performed with 10% v/v formamide. All data are presented as mean ± s.d. of ≥ 3 independent experiments. iSiMREPS showed a linear dynamic range of approximately 3 fM - 5 pM which is approximately 3.2 orders of magnitude.

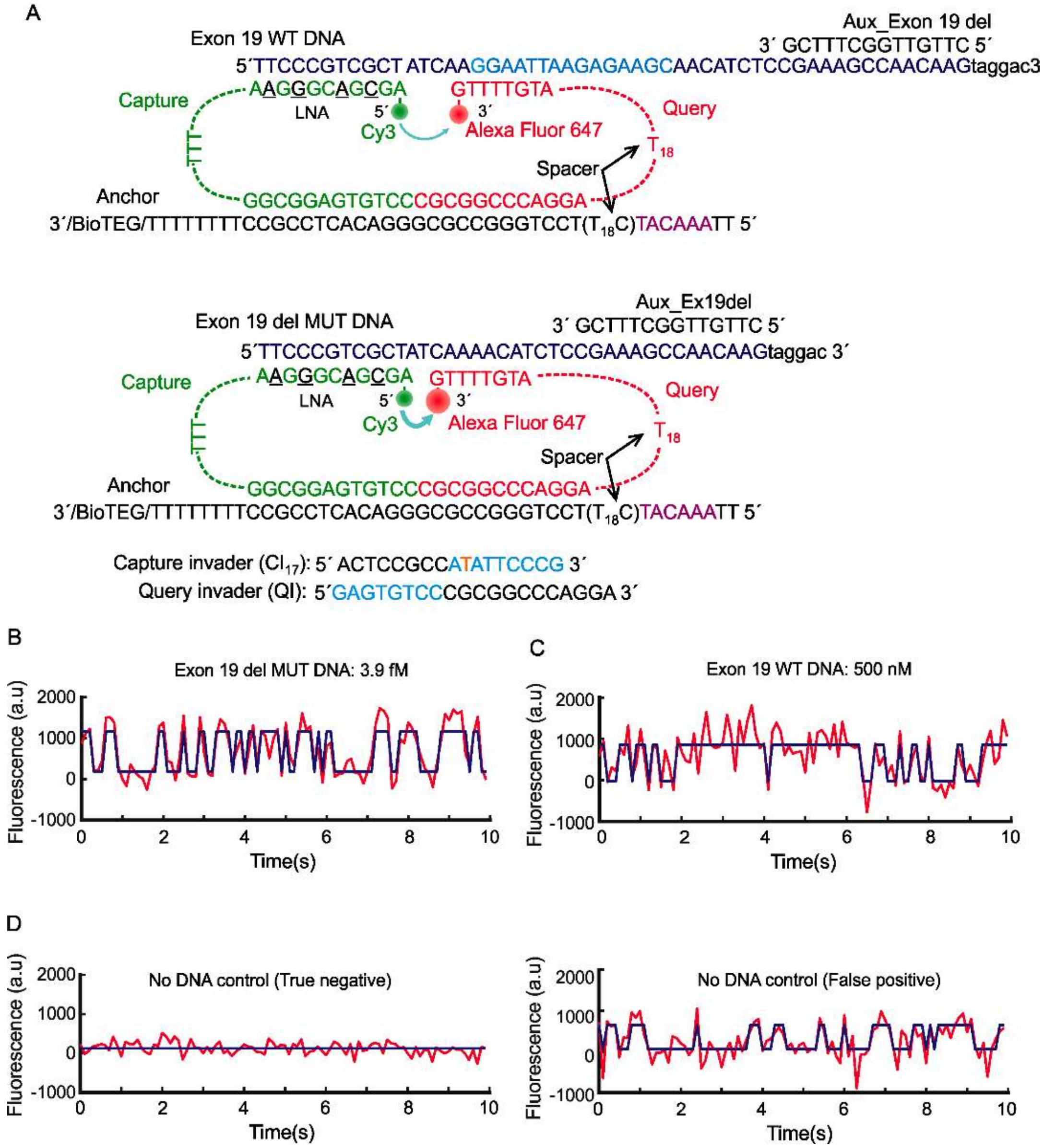


**Figure S14**. **Schematic of iSiMREPS sensor design for detecting *EGFR* exon 19 deletion mutant and wild type DNA and representative single molecule kinetic traces.** (A) Schematic of the iSiMREPS sensor for detection of *EGFR* exon 19 deletion mutant and wild-type DNA. The query probe was designed to be fully complementary to a short segment of the mutant DNA while lacking perfect complementarity with wild-type DNA. Representative single molecule kinetic traces for exon 19 deletion mutant DNA at 3.9 fM (B) and exon 19 wild type DNA at 500 nM (C). (D) Representative true negative and false positive single molecule kinetic traces for no DNA target (control).

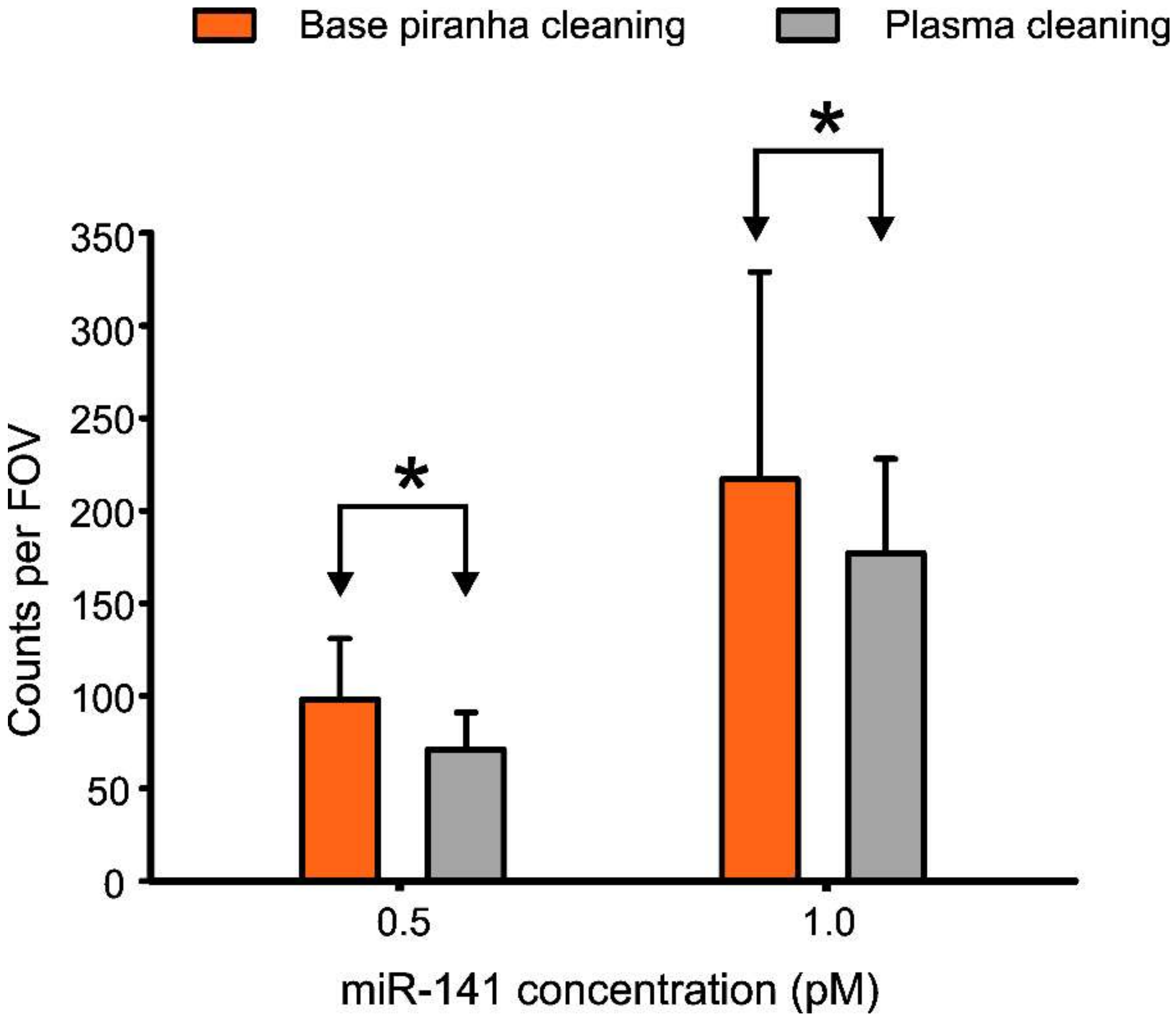


**Figure S15. Comparison of the performance of coverslip cleaning protocols for detecting miR-141.** Base piranha cleaning protocol used a solution consisting of 14.3% v/v of 28-30 wt% $NH_4OH$, and 14.3% v/v of 30-35 wt% $H_2O_2$ that was heated to 70-80°C, whereas plasma cleaning protocol used application of plasma for 3 min to clean glass coverslip. The experiments were performed using a glass coverslip passivated with biotin-PEG: m-PEG at a ratio of 1:100, 10 nM sensor, 0.5 and 1.0 pM miR-141, and 2 µM invaders ($CI_{mis}$ + QI). All imaging was performed with 10% v/v formamide. All data are presented as mean ± s.d. where n = 3 independent experiments. Single asterisk indicates the statistically insignificant differences at 95% confidence levels as assessed using a two-tailed, unpaired t test.
